# Genomic adaptations to an endolithic lifestyle in the coral-associated alga *Ostreobium*

**DOI:** 10.1101/2020.07.21.211367

**Authors:** Cintia Iha, Katherine E. Dougan, Javier A. Varela, Viridiana Avila, Christopher J. Jackson, Kenny A. Bogaert, Yibi Chen, Louise M. Judd, Ryan Wick, Kathryn E. Holt, Marisa M. Pasella, Francesco Ricci, Sonja I. Repetti, Mónica Medina, Vanessa R. Marcelino, Cheong Xin Chan, Heroen Verbruggen

## Abstract

The green alga *Ostreobium* is an important coral holobiont member, playing key roles in skeletal decalcification and providing photosynthate to bleached corals that have lost their dinoflagellate endosymbionts. *Ostreobium* lives in the coral’s skeleton, a low-light environment with variable pH and O□ availability. We present the *Ostreobium* nuclear genome and a metatranscriptomic analysis of healthy and bleached corals to improve our understanding of *Ostreobium*’s adaptations to its extreme environment and its roles as a coral holobiont member. The *Ostreobium* genome has 10,663 predicted protein-coding genes and shows adaptations for life in low and variable light conditions and other stressors in the endolithic environment. This alga presents a rich repertoire of light-harvesting complex proteins but lacks many genes for photoprotection and photoreceptors. It also has a large arsenal of genes for oxidative stress response. An expansion of extracellular peptidases suggests that *Ostreobium* may supplement its energy needs by feeding on the organic skeletal matrix, and a diverse set of fermentation pathways allow it to live in the anoxic skeleton at night. *Ostreobium* depends on other holobiont members for vitamin B12, and our metatranscriptomes identify potential bacterial sources. Metatranscriptomes showed *Ostreobium* becoming a dominant agent of photosynthesis in bleached corals and provided evidence for variable responses among coral samples and different *Ostreobium* genotypes. Our work provides a comprehensive understanding of the adaptations of *Ostreobium* to its extreme environment and an important genomic resource to improve our comprehension of coral holobiont resilience, bleaching and recovery.

## Introduction

Coral health depends on the harmonious association between the coral animal and its microbial associates, together known as the holobiont. A wealth of studies shows that climate change threatens coral health by disrupting the association between the coral and its photosynthetic dinoflagellate endosymbionts (Symbiodiniaceae), culminating in coral bleaching and death. The role of other microbes in coral resilience is just starting to be understood, and our current knowledge is largely based on correlations between coral health status and the presence of microbial taxa inferred from metabarcoding. Whole-genome sequences of corals and their symbionts are an invaluable resource to obtain a mechanistic understanding of the functioning and resilience of the holobiont. Recent genomic studies have shown that the coral host, dinoflagellate symbionts, and prokaryotes living in the coral tissue have complementary pathways for nutrient exchange, highlighting the interdependence and possible co-evolution between these members of the coral holobiont [1, 2]. To date, most work has focused on the coral animal and microbiota associated with its living tissue, with very little work done on the highly biodiverse and functionally important microbiota inhabiting the skeleton of the coral [3, 4].

The green alga *Ostreobium* sp. is an important eukaryotic symbiont living inside the coral skeleton (Figure 1A) [4, 5]. This endolithic alga is the principal agent of coral reef bioerosion, burrowing through the limestone skeleton of corals and other reef structures and dissolving up to a kilogram of reef CaCO_3_ per m^2^ per year [6]. These algae bloom in the skeleton when corals bleach (Figure 1B) [7, 8], boosted by the extra light, and – hypothetically – the extra nitrogen and CO_2_ that may reach the skeleton in the absence of Symbiodiniaceae. Part of the carbohydrates produced by *Ostreobium* photosynthesis makes their way into the coral tissue, extending the time it can survive without its dinoflagellate partner [9, 10]. While these ecological and physiological phenomena have been described, our knowledge of the molecular mechanisms involved is scarce, severely limiting our understanding of key processes in healthy and bleached holobionts. The question remains as to not only what is the mechanism of this skeletal deterioration, but how tightly *Ostreobium* metabolism integrates with that of the coral and associated microbiota. Developing this knowledge will be essential to understand and manage the roles that skeletal microbiota play during coral bleaching.

**Figure 1.**
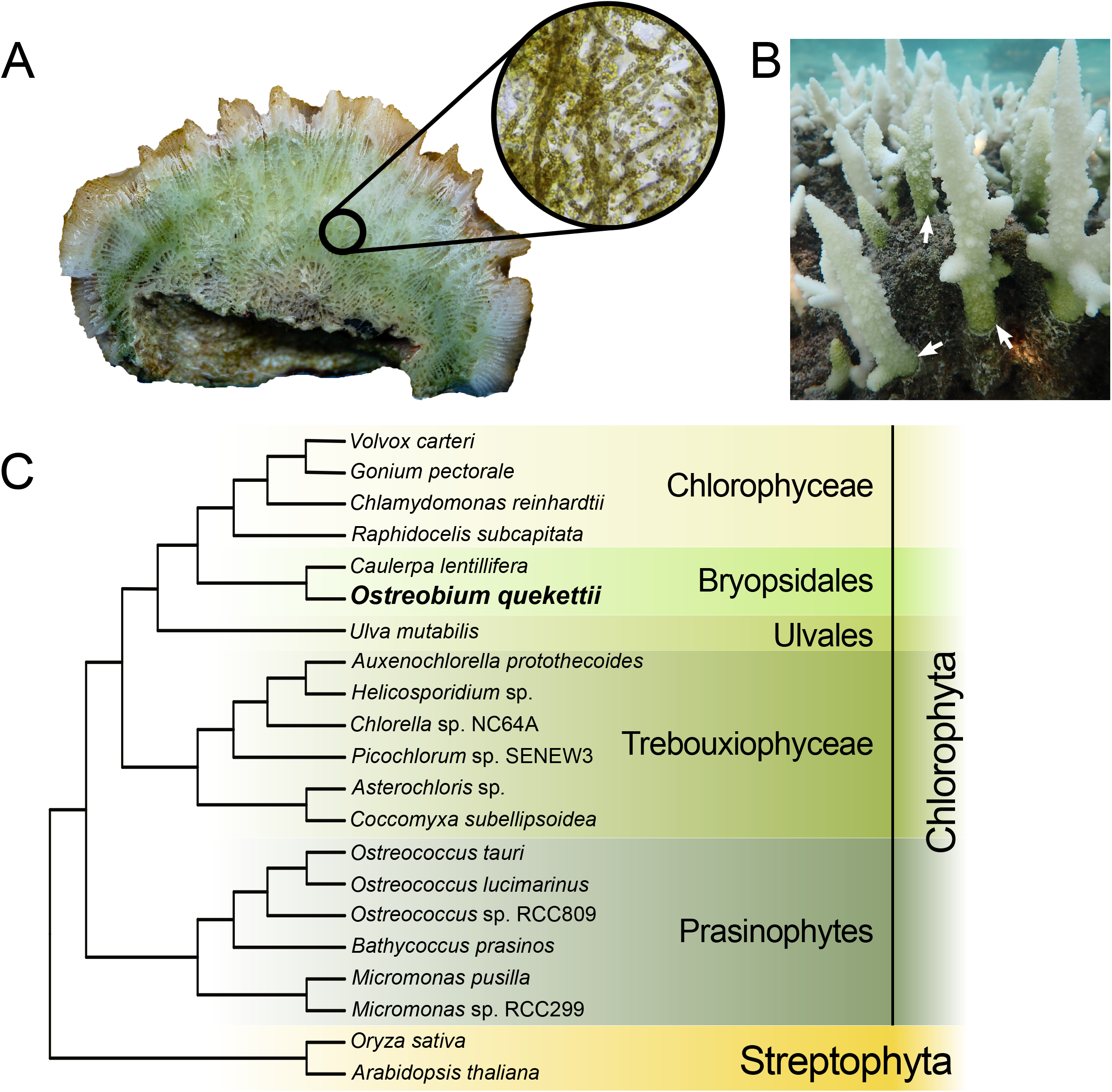
Localisation of *Ostreobium* in the coral skeleton and phylogenetic position. (A) Cross-section of *Paragoniastrea australensis* coral showing *Ostreobium* that inhabits the skeleton. The inset shows *Ostreobium* filaments after skeletal decalcification. (B) Bleached coral with evident *Ostreobium* bloom (indicated by white arrows). Photograph by Alexander Fordyce. (C) Phylogenetic tree of Viridiplantae (Streptophyta + Chlorophyta) showing the position of *Ostreobium quekettii*.

*Ostreobium* has an extreme lifestyle for a green alga [11]. It lives in a very dimly lit environment, with mostly the low-energy, far-red wavelengths not used up by Symbiodiniaceae available [12]. The action spectrum of *Ostreobium* photosynthesis extends into far-red wavelengths, but the underlying molecular mechanisms are little known [13, 14]. The daily rhythm of oxygenic photosynthesis and respiration within the skeletal matrix leads to strong fluctuations of pH and O_2_, ranging from total anoxia at night to ca. 60% of air saturation during the day [11, 15], and the skeleton limits diffusion of O_2_ and other compounds. Free-living algae do not normally encounter these stressful conditions, and the mechanisms allowing *Ostreobium* to thrive in this extreme habitat are virtually unknown.

From an evolutionary perspective, *Ostreobium* is a member of the green plant lineage (Viridiplantae), which includes the land plants that originated in the Streptophyta lineage and a broad diversity of algae in the Chlorophyta lineage [16] (Figure 1C). *Ostreobium* is in Bryopsidales, an order of marine algae that has evolved in the Chlorophyta. While *Ostreobium* forms microscopic filaments, many representatives of this order are larger seaweeds [17].

So far, the genomic resources available for coral holobiont research have been limited to the coral animal, its dinoflagellate photosymbionts, and a small fraction of the prokaryotes associated with its tissue (e.g. [1] [2] [18] [19] [20]). Here, we present the first nuclear genome of an *Ostreobium* species to extend the available genomic toolkit into the coral skeleton, an element of the holobiont crucial for our comprehension of coral resilience, bleaching, and recovery. We expect that these genomic resources will spur new insights into processes of coral bleaching encompassing the entire holobiont, as this knowledge will be essential to safeguard the future of coral reefs in a changing climate. Based on comparative analyses of the *Ostreobium* genome with those of other green algae, we show *Ostreobium*’s innovations in light-harvesting antennae and derive insights and hypotheses about its functions as a coral symbiont, and as an alga living in an extreme environment.

## Results and Discussion

We obtained a draft nuclear genome of *Ostreobium quekettii* (SAG culture collection strain 6.99, non-axenic) by assembling sequence reads from Illumina and Nanopore platforms (Data S1). We assembled the final haploid nuclear genome in 2,857 scaffolds (total 146.26 MB of assembled bases; N50 length 73.43 KB), with an average sequence coverage of 99.72x for short-read data and 25.2x for long-read data. The genome is diploid with ~1.29% heterozygosity. The GC content is 52.4%, which is higher than 40.4% of *Caulerpa lentillifera* (the closest relative of *Ostreobium* for which a nuclear genome has been sequenced), but less than most other green algae (Data S1). Gene annotation of the haploid representation of the genome resulted in 10,663 predicted protein-coding genes, of which RNA sequencing data supported ~52%. We recovered 60.7% of the complete core conserved BUSCO eukaryote genes, which is similar to *C.lentillifera* (59.2%) and *Ulva mutabilis* (65.8%) (Data S1).

### Photobiology in a dark place

Our work shows that *Ostreobium* has more light-harvesting complex (LHC) proteins than most green algae and *Arabidopsis thaliana* (Figure 2A, Figure S1). This expansion of the LHC protein arsenal is found in both photosystems, with duplications of the *Lhca1* and *Lhca6* gene families associated with PSI (Figure S1A) and the presence of both *Lhcp* and *Lhcb* families associated with PSII (Figure S1B).

**Figure 2.**
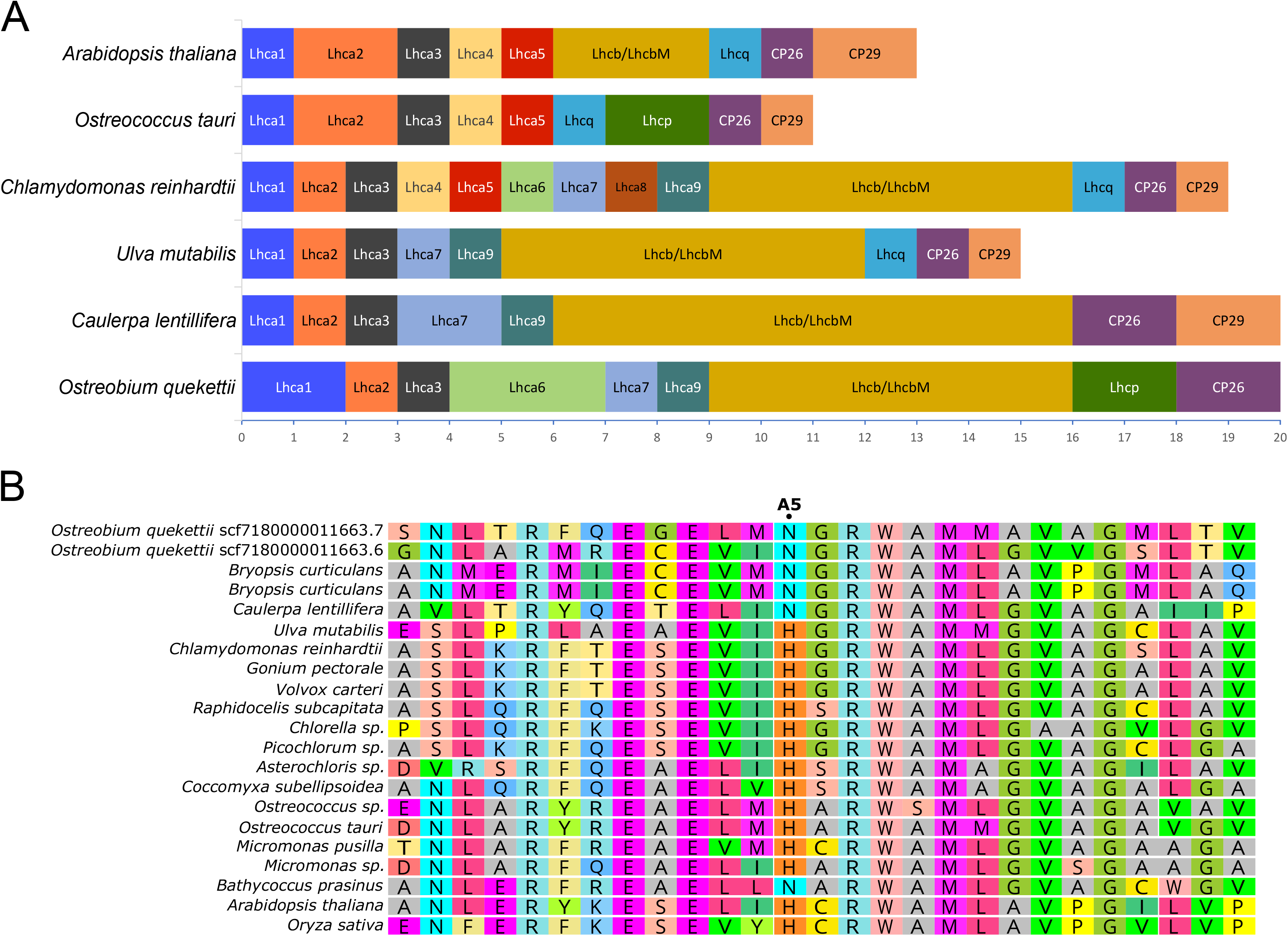
Diversity of light-harvesting complex proteins in *Ostreobium* and some other species. **(A)** Number of different light-harvesting complex (LHC) proteins in *Ostreobium*, other green algae and *Arabidopsis thaliana.* For a comparative analysis, we included the species models for plants and green algae, *Arabidopsis thaliana* and *Chlamydomonas reinhardtii*, respectively; *Ostreococcus tauri,* because it presents a special type of LHC protein that are found mainly in Prasinophytes (Lhcp) and were previously well characterised [25]; and *Ulva mutabilis* and *Caulerpa lentillifera* because they are ulvophycean relatives of *Ostreobium*. (B) Amino acid sequence comparison between Lhca1 proteins, showing asparagine (N) at the chlorophyll-binding residue A5 in *Ostreobium*. See also Figure S1.

Most green algae possess a single *Lhca1* gene, while *Ostreobium* has two *Lhca1* (Figure 2). The amino acid residue binding chlorophyll in LHC proteins is essential in determining the chromophore organisation, which affects light spectral absorption [21]. The Lhca1 protein typically uses histidine as the chlorophyll-binding residue (A5 site in Figure 2B), but the siphonous green algae (*Ostreobium*, *C. lentillifera* and *Bryopsis corticulans*) all have asparagine (Figure 2B). *Arabidopsis thaliana* mutants with asparagine at the A5 site were shown to have red-shifted absorption spectra [21, 22], suggesting that siphonous green algae may also use this mechanism to access far-red wavelengths for photosynthesis. The Lhca6 protein, located in the outer LHC belt, forms heterodimers with Lhca5, and their long C-terminal loops facilitate interactions between the inner and outer LHC belts [23, 24]. While most microalgae have a single *Lhca6* copy (and none in the seaweeds *C. lentillifera* and *U. mutabilis*)*, Ostreobium* contains three copies (Figure S1A), supported by different chromosomal contexts and their presence in our transcriptome.

In the PSII-associated LHC, *Ostreobium* possesses an unusual combination of both the Lhcp and major light-harvesting complex (Lhcb) protein families (Figure 2A and Figure S1B). Lhcb is found in most species of the green lineage except prasinophytes that use the Lhcp family instead [25]. Only the streptophyte *Mesostigma viridis* is known to encode proteins of both families (Figure S1B), suggesting that Lhcp and Lhcb were both part of the LHCII antenna system of the green plant ancestor and the families were differentially lost in different green algal and land plant lineages [26, 27].

Although *Ostreobium* shows a high diversity of LHC genes, it lacks many genes for photoprotection and photoreceptors. The non-photochemical quenching (NPQ) genes for LHCSR and PsbS are both absent from *Ostreobium* and *C. lentillifera* genomes (Figure S1B). Energy-dependent quenching (qE) was not observed in most siphonous green algae [28], and our genomic data lack crucial genes for this process. *Ostreobium* also lacks all the light-harvesting complex-like (LIL) genes coding for OHP1, OHP2, LIL3, ELIP, and four-helix proteins. While the function of LIL proteins has not been comprehensively determined, their involvement in response to light stress is known [29]. The loss of genes involved in high-light sensitivity also extends to the chloroplast genome of *Ostreobium,* which lacks the chloroplast envelope membrane protein gene (*cem*A) that is needed in *Chlamydomonas* to persist in high-light conditions [30].

*Ostreobium* has fewer known photoreceptors than do most other green algae. We identified three blue light photoreceptors: a phototropin (not shown), a plant cryptochrome, and a photolyase/blue-light receptor (PHR1; Figure S2). Given the predominance of far-red light in coral skeletons, one might expect the red/far-red phytochromes to be present in *Ostreobium*, but this was not the case. Although phytochromes are widely found in the green plant lineage, they have been lost in most of the green algal lineage Chlorophyta [31, 32]. Most green algae appear to have no specific photoreceptors for red light [33], but the animal-like cryptochrome in *Chlamydomonas* can be activated by red light [34]. This protein’s functions include the transcription of genes involved in photosynthesis, pigment biosynthesis, cell cycle control and circadian clock [35]. This gene is not found in *Ostreobium*, which also lacks the Cry-DASH-type cryptochromes observed in many other green algae (Figure S2). *Ostreobium* does have two genes similar to the *Arabidopsis* putative blue-light receptor protein called PAS/LOV (Uniprot O64511) [36]. *Ostreobium* also lacks the rhodopsin-like photoreceptors.

Besides the photoprotective LHC genes and photoreceptors, *Ostreobium* and *C. lentillifera* appear to have lost the light-dependent protochlorophyllide oxidoreductase (LPOR), another important light signalling gene involved in chlorophyll biosynthesis. The reduction of protochlorophyllide to chlorophyllide can be catalysed by either of two non-homologous enzymes: the nuclear encoded LPOR and the plastid-encoded light-independent (DPOR) protochlorophyllide oxidoreductase [37]. Most green algae have both systems, and DPOR has been lost in many eukaryotes [38]. The *Ostreobium* genome provides the first evidence for the loss of LPOR in any eukaryote, and a screening of *C. lentillifera* also came back negative, suggesting that the loss may have occurred in the common ancestor of Bryopsidales. Both genera encode DPOR in their chloroplast genomes [30, 39]. Although both enzymes catalyse the same reaction, they have different features: DPOR is encoded, synthesised and active in the plastid, and is highly sensitive to oxygen [40], while the nucleus-encoded LPOR is synthesised in the cytosol and active in the plastid, and requires light to be activated [41]. DPOR might be an advantage over LPOR for endolithic photosynthetic organisms because of the low-light, low-oxygen environments they inhabit [4].

The *Ostreobium* genome clearly reflects its evolutionary trajectory into a peculiar light habitat, with an unparalleled arsenal of LHC proteins but few known mechanisms to sense the light or protect itself against excessive light. Some of these genome features are shared with other Bryopsidales, including the loss of LPOR and qE-type non-photochemical quenching. This suggests that the common ancestor of Bryopsidales may have been a low-light-adapted organism, possibly an endolithic alga like *Ostreobium* is now, a hypothesis supported by *Ostreobium* being the sister lineage of all other Bryopsidales [42] and other bryopsidalean lineages also containing old, largely endolithic families [43]. Bryopsidales originated in the Neoproterozoic with *Ostreobium* diverging in the early Paleozoic [17, 44]. One could speculate that the bryopsidalean ancestor inhabited a low-light environment, possibly on the dimly lit seafloor beneath Cryogenian ice sheets. The different lineages emerging from this ancestor could then have followed different evolutionary trajectories during the onset of Paleozoic grazing, with the *Ostreobium* lineage fully committing to an endolithic lifestyle while other bryopsidalean lineages engaged in an evolutionary arms race with grazers to form larger and chemically defended macroalgae.

The unusually large arsenal of LHC proteins appears to be confined to the *Ostreobium* lineage. The genus is present in a diverse range of light environments, from old oyster shells in the intertidal, where it experiences full-spectrum sunlight, to the coral skeleton (far-red enriched) and in mesophotic habitats (blue light), hinting at its capability to harvest energy from photons of many different wavelengths [45]. While from the genome data alone, we cannot derive that the different LHC proteins convey a capability for photosynthesis at different wavelengths, it is known that different molecular arrangements of LHC proteins and the specific pigments binding to them have an impact on their spectral properties [46], which could help *Ostreobium* acclimate to different light environments.

### Life in an extreme environment

The endolithic environment, and the coral skeleton, is an extreme environment in many ways. Oxygen levels vary strongly, from high concentrations caused by photosynthesis during the day to complete anoxia due to respiration during the night, and this trend is mirrored in strong diurnal pH fluctuations [4, 15]. Reactive oxygen species (ROS) can be produced in high quantities in these conditions, particularly in the morning when photosynthesis starts [47].

The *Ostreobium* genome exhibits a strong genetic capacity for oxidative stress response, with ROS scavenging and neutralising genes present in large numbers compared to other green algae (Figure 3). We found five copies of catalase (CAT), the enzyme that processes hydrogen peroxide. Most other green algae have one or two (Figure 3), or none in the case of prasinophytes (Figure S3). Four of the *Ostreobium* catalases formed a unique lineage in our phylogenetic analysis (Figure S3), and three of them are found in tandem in the same scaffold, indicating diversification of this gene in the *Ostreobium* lineage. Hydrogen peroxide can also be processed through the glutathione-ascorbate cycle, a metabolic pathway neutralising hydrogen peroxide through successive oxidations and reductions of ascorbate, glutathione, and NADPH [48] (Figure 3). *Ostreobium* featured high copy numbers of the enzymes to quickly purify ascorbate, with five copies of ascorbate peroxidase (APX) and four monodehydroascorbate reductases (MDHAR) (Figure S3). The glutathione-ascorbate cycle is important to keep ascorbate free for H_2_O_2_ scavenging (Figure 3).

**Figure 3.**
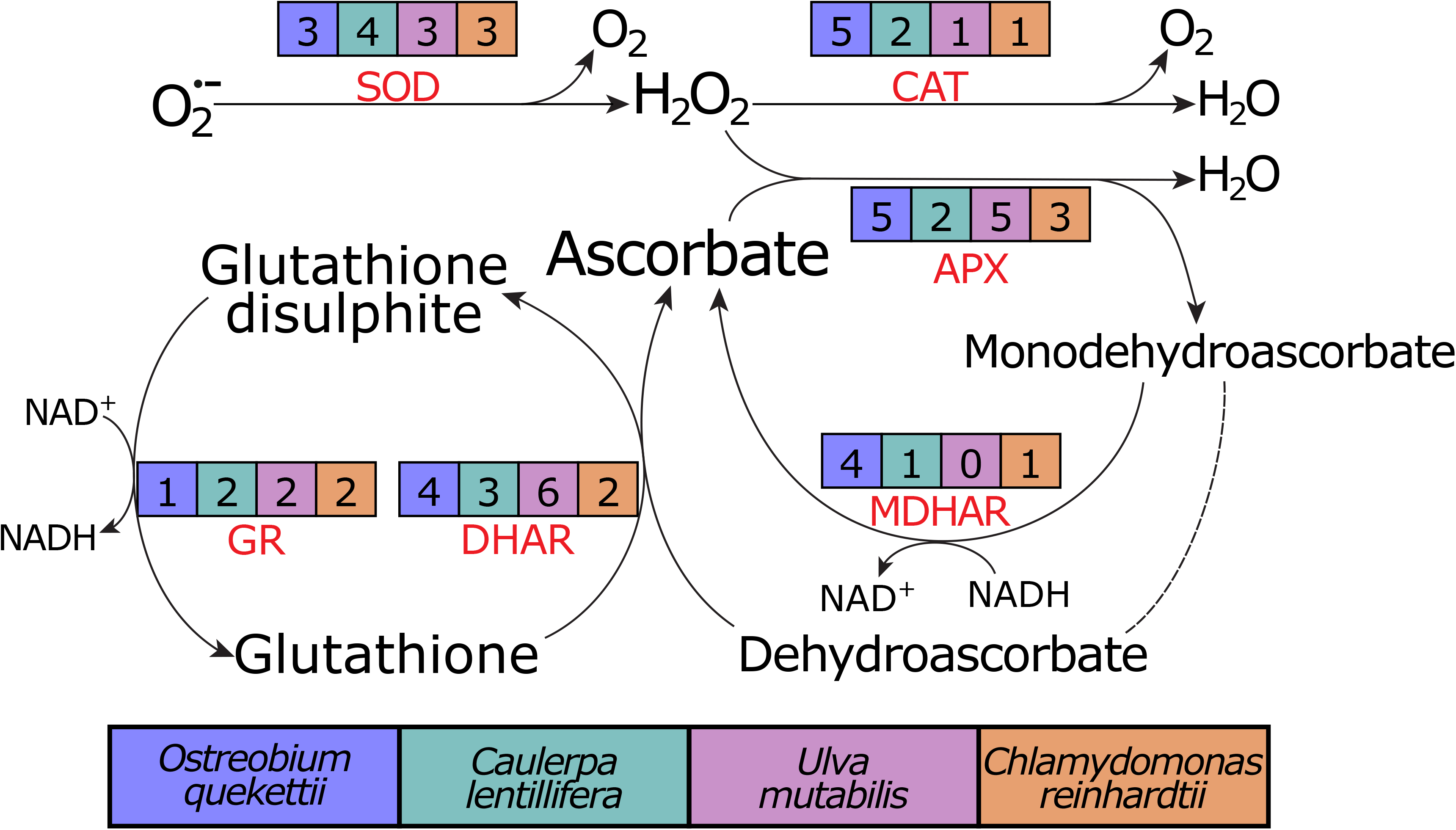
Simplified oxidative response pathway comparing the number of genes for enzymes found in the genomes from *Ostreobium* and some other green algae. Compared with *C. lentillifera*, *U. mutabilis* and C*. reinhardtii*, *Ostreobium* does have more copies of genes related to quick response to neutralise ROS, such as catalase (CAT) and monodehydroascorbate reductase (MDHAR). SOD, superoxide dismutase; APX, ascorbate peroxidase; DHAR, dehydroascorbate reductase; GR, glutathione reductase. See also Figure S3.

Most algae live in environments with higher oxygen concentrations and can produce energy via the respiratory electron transport chain. In the coral skeleton, however, any oxygen produced through photosynthesis is completely consumed by respiration within an hour of the onset of darkness [15], and the environment is anoxic for long periods. *Ostreobium* does have a variety of options for fermentative metabolism (Figure S4), including expanded copy numbers for LDH, ALDH and MDH (Data S1), implying that acetate, succinate and lactate may be the preferred fermentation products in *Ostreobium*.

A comparison of the number of genes having particular InterPro annotations between *Ostreobium* and *Caulerpa* showed a large number of depleted IPR terms in *Ostreobium*, which can be attributed to an enrichment in the *Caulerpa* genome, as counts in *Ostreobium* are comparable to those of other green algae. However, peptidases were strongly enriched in *Ostreobium* (e.g. Peptidase S1, PA clan – 189 genes; Serine-proteases trypsin domain – 159 genes; Figure S5). About 40% of the genes are predicted to be secreted (signal peptide), and 25% membrane-bound, compared to 31% and 12% (of total 57 genes) respectively in *C. lentillifera*, suggesting an expanded potential for external protein degradation in *Ostreobium*. It is known that *Ostreobium* can penetrate the organic-inorganic composite material of the black pearl oyster nacreous layer [49], and the combination of proteolytic enzyme proliferation and its growth at excessively low light intensities lends support to the hypothesis that this alga could complement its energy needs by feeding on the organic matrix of the coral skeleton and shells.

### The coral holobiont

*Ostreobium* plays a number of important roles in the coral holobiont, particularly during periods of coral bleaching [4], but current knowledge is far from complete, and the genome can help define hypotheses of how the species may interact with other holobiont members.

#### Molecular mechanism of CaCO_3_ dissolution

*Ostreobium* and endolithic fungi play important roles as microbial bioerosion agents in the coral skeleton [50]. While the molecular mechanisms behind this phenomenon in eukaryotes are not yet known [51], the genome allows us to make conjectures about how bioerosion by *Ostreobium* might occur (Figure 4A). A working model for microbial carbonate excavation was first described for cyanobacteria, involving passive uptake of Ca^2+^ at the boring front, decreasing the ion concentration in the extracellular area below calcite saturation levels and leading to the dissolution of adjacent calcium carbonate [52]. Imported Ca^2+^ is transported along the cyanobacterial filament and excreted away from the growing tip, likely by P-type Ca^2+^-ATPases that pair transport of Ca^2+^ with counter transport of protons [52].

**Figure 4.**
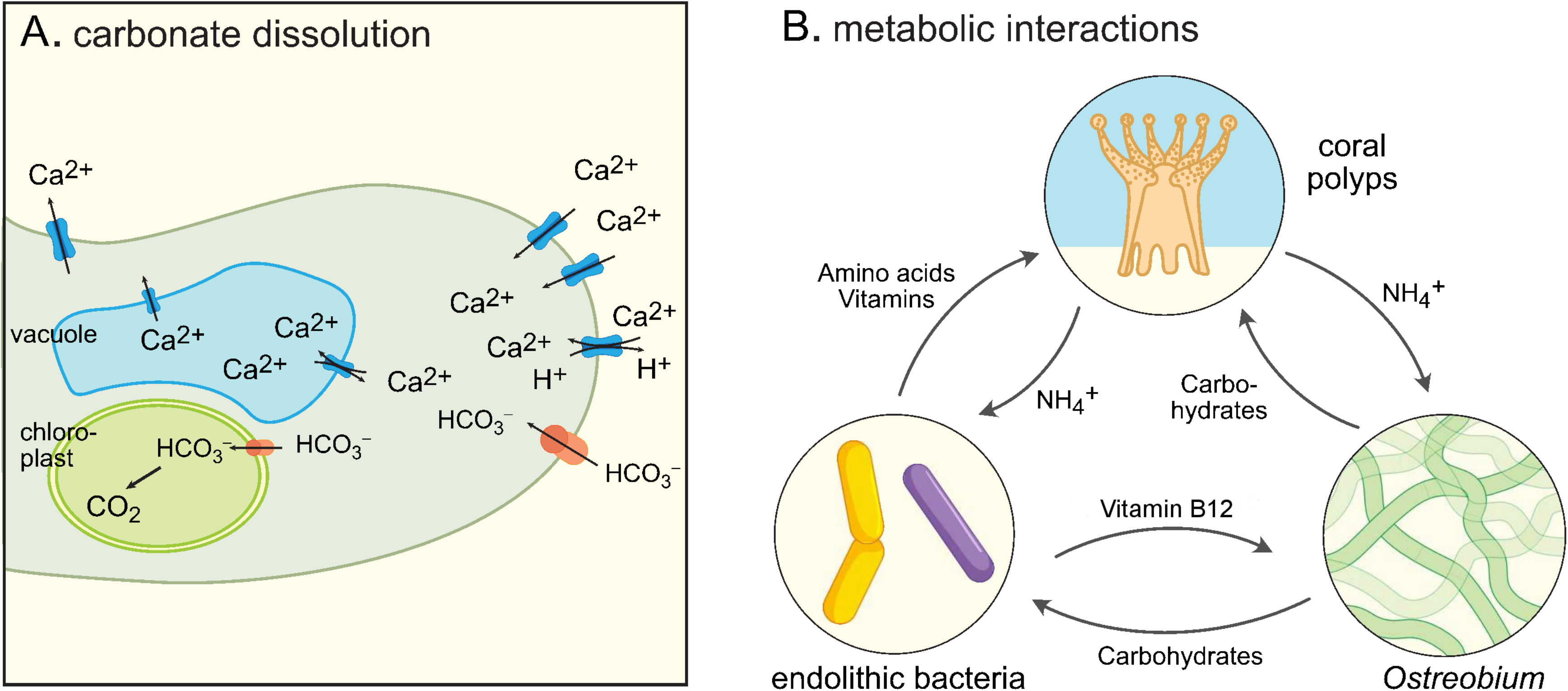
Roles of *Ostreobium* in the coral holobiont. (A) Potential mechanisms available to *Ostreobium* for excavation of the CaCO_3_ skeleton of corals near the growing tip of *Ostreobium* filament. (B) Possible interactions between members of the holobiont derived from genome sequence data.

*Ostreobium* has an expanded repertoire (34 genes) of calcium transporters (Data S1), including 19 voltage-dependent calcium channels and several transient receptor potential transporters, two-pore channels and calcium-transporting ATPases. Calcium uptake, possibly combined with acidification via a Ca^2+^/H^+^ transporter, would promote decalcification (Figure 4A), allowing *Ostreobium* to burrow into the coral skeleton. Calcium toxicity can be avoided either by accumulation in a vacuole and/or posterior transport out of the cell. In addition to calcium transport, bicarbonate uptake could also play a role in the burrowing mechanism. *Ostreobium* carries two orthologs of the CIA8 transporter responsible for bicarbonate transport in *C. reinhardtii* [53], and the imported bicarbonate may be further transported into the chloroplast to be fixed. *Ostreobium* has a carbonic anhydrase predicted to be targeted outside the cell that may further assist the decalcification process. In natural communities dominated by *Ostreobium*, CaCO_3_ dissolution was higher at night, suggesting that *Ostreobium* takes advantage of the lower external pH at night [51].

Even though the *Ostreobium* genome does not suggest a specific mechanism, it lends support to the notion that its bioerosion likely bears similarities to the process in cyanobacteria. A detailed characterisation of this process should be a priority, as *Ostreobium* is responsible for ca. 30-90% of microbial dissolution of skeletal CaCO_3_. Higher temperatures and lower pH boost this activity, suggesting that this yet unknown process will lead to major reef deterioration in future ocean conditions [51].

#### Interactions with other holobiont members

As far as holobiont functioning goes, interactions between the coral animal and its algal and bacterial symbionts are best understood, and interactions between the Symbiodiniaceae and bacteria are just starting to come into focus [54], but little is known about interactions involving *Ostreobium*. It is well known that several algae are auxotrophic for certain nutrients, e.g. vitamin B12 (cobalamin), a cofactor involved in the synthesis of methionine that many algae obtain from associated bacteria [55]. The *Ostreobium* genome shows that the metabolic pathways involved in the production of vitamins B1, B2, B6 and B9 are complete, but it is auxotrophic for vitamin B12 (cobalamin). The *Ostreobium* genome encodes a B12-independent methionine synthase (METE) in addition to the B12-dependent version (METH, [56]), so while the alga most likely does not strictly require B12 for growth, the presence of METH suggests that it uses B12 provided by other holobiont members. Corals are also auxotrophic for several vitamins and amino acids that are produced by holobiont members [2] (Figure 4B).

The nature of metabolic exchanges between *Ostreobium* and the coral animal is an open question in holobiont research, and a potentially critical one, as coral bleaching and subsequent blooming of the endolithic *Ostreobium* algae become increasingly common due to ocean warming. Typical algal-animal metabolic exchanges include nitrogen and CO_2_ provision by the animal to the alga and carbohydrate provision to the animal by the alga [57, 58]. Coral polyps are known to secrete nitrogen in the form of ammonia. An expanded repertoire of ammonia transporters was identified in *Ostreobium* (Data S1), potentially reflecting an adaptation to increase and diversify ammonia uptake in the alga. This observation is also in line with the presence of diazotrophic bacteria facilitating the conversion of N_2_ into ammonia in marine limestones (including coral skeletons) and ammonia being the most abundant form of inorganic nitrogen in skeletal pore waters [59]. It adds to the evidence for the roles that endolithic organisms play in the holobiont N cycle. During carbon cycling in the holobiont, glucose has been postulated as one of the main carbohydrates exchanged between Symbiodiniaceae and corals [57]. While carbon compounds fixed by endoliths are known to be transferred to the coral animal and subsequently assimilated, neither the exact transferred molecules nor the molecular mechanisms involved in their translocation have been characterised, but the *Ostreobium* genome encodes four genes coding for H^+^-glucose transporters that might be involved in this process (Data S1).

#### Probing changes in holobiont processes

While these investigations of the *Ostreobium* genome allowed us to evaluate hypotheses about interactions within the holobiont, gaining more in-depth insight will require approaches that study multiple partners simultaneously. As a first step towards understanding the molecular mechanisms in the coral holobiont during coral bleaching, we screened transcriptomes of healthy and bleached coral holobionts. We exposed fragments of the coral *Orbicella faveolata* to elevated temperatures leading to bleaching, followed by re-acclimation of the bleached samples to ambient temperatures (post-bleaching condition), during which the corals remained bleached, and an *Ostreobium* bloom occurred. Total metatranscriptomes (including coral tissue and skeleton) were generated for healthy control samples that were kept under ambient temperature (no bleaching), and the bleached samples.

Taxonomic profiles of the reads and assembled transcripts are provided in Data S1. While the metatranscriptome sequencing depth was not designed to track up-and down-regulation of individual genes, it was clear that expression of the photosynthesis genes *psb*A and *rbc*L from Symbiodiniaceae was drastically lower in bleached samples while expression of these genes in *Ostreobium* increased (Figure 5A), supporting the notion that *Ostreobium* becomes a dominant agent of photosynthesis in the bleached holobiont. The bleached state increases the light available to *Ostreobium*, which likely leads to the need for higher repair rates of PSII protein D1 (encoded by *psb*A) and higher rates of carbon fixation (facilitated by RuBisCO, encoded by *rbc*L). The expression of these genes by Symbiodiniaceae in the bleached samples, albeit at low levels, suggests that some of these endosymbionts have remained or that some re-colonisation has occurred during the post-bleaching period.

**Figure 5.**
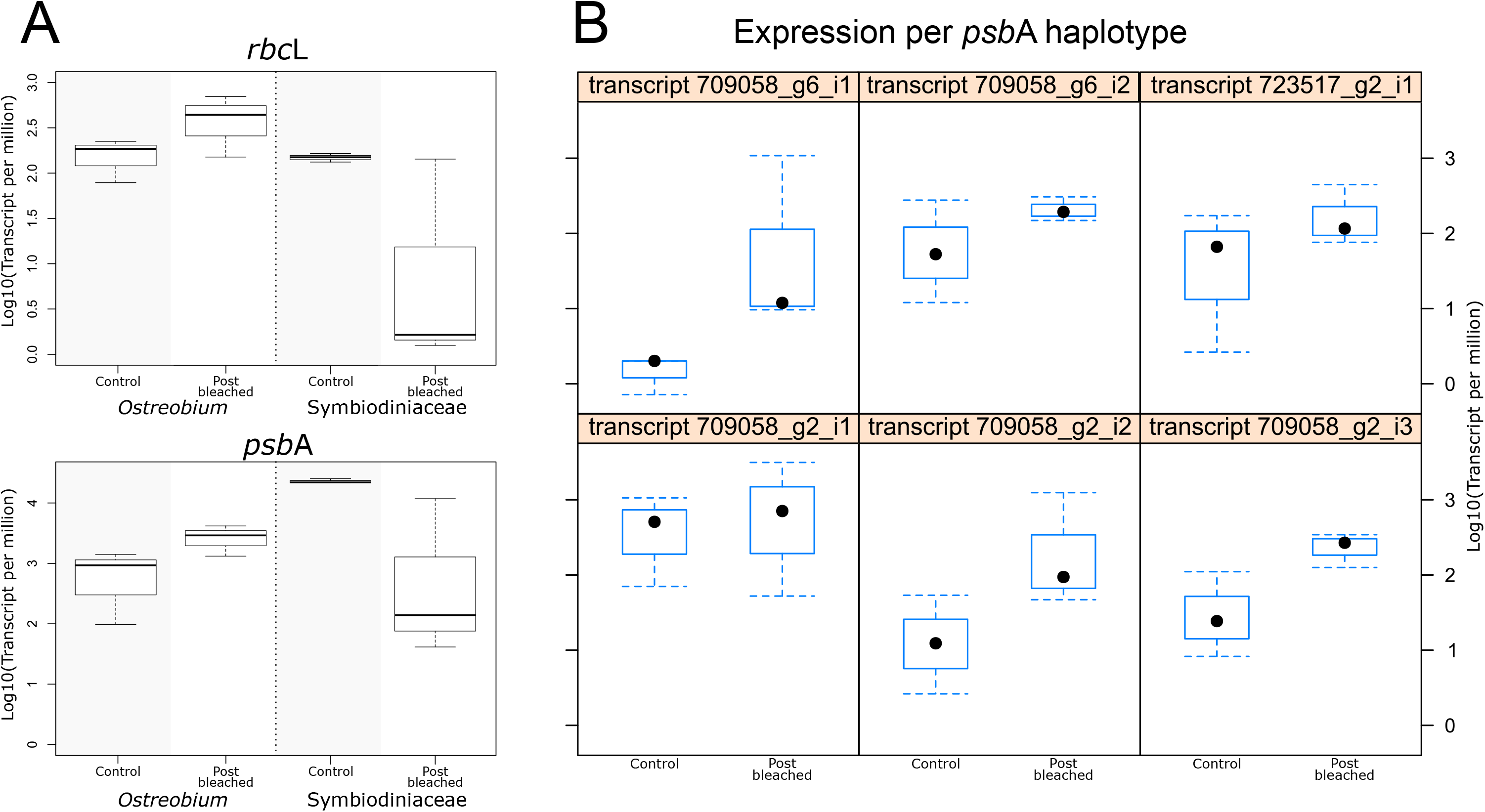
Individual gene expression in the metatranscriptome analysis. (A) Comparison of *rbcL* and *psbA* gene expression between *Ostreobium* combined transcripts (haplotypes) and Symbiodiniaceae transcripts in control and post-bleached treatments. (B) Expression between control and post-bleached treatments of individual *psbA* haplotypes found in *Ostrebium* transcriptomes, indicating several strains in *O. faveolata* skeletons.

We detected multiple haplotypes of *psb*A (Figure 5B), indicating that several strains of *Ostreobium* were present in the *O. faveolata* skeletons [60]. While expression levels for these haplotypes tended to increase, some did not change significantly while others differed by an order of magnitude or more. These differences suggest that *Ostreobium* strains may differ physiologically or change in relative abundance during the experiment. Such differences in the microbiome – whether *Ostreobium* strains or other holobiont members – may result in high variability among coral samples in experimental work. Indeed, we found considerable differences in expression levels between samples within conditions, suggesting that future metatranscriptome experiments should be planned to use generous replication. The metatranscriptomes also provide some hints as to where *Ostreobium* may source its needs for vitamin B12 (cobalamin). We detected 29 bacterial transcripts from the vitamin B12 pathway, including several Proteobacteria, Bacteroidetes, and Cyanobacteria that are known to be abundant in the coral skeleton (Data S1) [43]. These results contribute to defining potential mutualistic relationships between bacteria and *Ostreobium* in the holobiont (Figure 4B).

## Conclusion

The complexity of the coral holobiont presents an interesting challenge to reconstruct a comprehensive model of metabolic exchanges and other interactions among its component organisms. Recent progress in building such models from genomes of coral and tissue-associated prokaryotes [2] has not been mirrored in the skeleton. Our results allowed us to refine hypotheses from previous physiological work, illuminating the biology of a keystone eukaryotic phototroph in this environment. Our work clearly shows genomic adaptations of *Ostreobium* to the low light and variable oxygen conditions it experiences in endolithic environments, with an unparalleled arsenal of light-harvesting complexes and expansions in pathways for fermentation and reactive oxygen processing. Despite this progress, many questions remain about the mechanisms involved in this alga’s interaction with the holobiont, their immediate physiological effects on the partners and longer-term ecological consequences. One challenge in this area of research is that the roles and impacts of *Ostreobium* vary in time, with a relatively minor contribution to holobiont photosynthesis in healthy corals [10]. However, our results are showing that *Ostreobium* blooms during bleaching cause a shift of expression levels between microbiome members, with *Ostreobium* becoming the dominant oxygenic phototroph. These changes likely have implications flowing through the entire microbiome interaction network, but our knowledge of this is in its infancy. Important questions remain as to what extent the *Ostreobium* bloom leads to a beneficial metabolite exchange with the coral host, and how the fitness costs/benefits of *Ostreobium* add up across the entire coral life cycle. The fact that there are >80 different species-level operational taxonomic units in *Ostreobium* complicates the matter further [43, 60]. Genome-scale data from *Ostreobium* and other interacting partners critically link these physiological features to underlying molecular mechanisms. The results and datasets generated in this study provide a foundational reference for future research into the biology of this key holobiont member, and the intricate role that *Ostreobium* plays in coral biology. This will be of particular importance in the light of global climate change, as the increased frequency of bleaching and lower pH will boost *Ostreobium* populations and carbonate dissolution rates.

## Acknowledgements

Funding was provided by the Australian Research Council (FT110100585 to H.V., DP150100705 to H.V. and C.X.C., DP190102474 to C.X.C., DP200101613 to H.V. and M.M.), the University of Melbourne (CBRI to H.V. & K.E.H.), the DoE Joint Genome Institute (CSP grant 1622 to M.M. and V.A.M.) and the National Science Foundation (OCE 1442206 and IOS 0644438 to M.M.), Pennsylvania State University (to M.M.), the Canon Foundation (to M.M.) and CONACyT (216837 to V.A.M.). This work is supported by the computational resources of the National Computational Infrastructure (NCI) National Facility systems through the NCI Merit Allocation Scheme (Project d85) awarded to C.X.C., and through the Nectar Research Cloud. We thank R. Iglesias-Prieto, C.T. Galindo and M. Weber for assisting with field experiments and J. Beardall, J.C. Lagarias and A.H. Knoll for discussions. We thank A. Fordyce for the bleached coral photograph.

## Author contribution

C.I., K.E.D., C.J.J., V.A., V.R.M., M.M., C.X.C., H.V. designed research; C.I., K.E.D., V.A., C.J.J., Y.C., L.M.J. performed research; K.E.H., H.V. contributed new reagents/analytic tools; C.I., K.E.D., J.A.V., V.A., C.J.J., K.A.B., Y.C., R.W., V.R.M., C.X.C., H.V. analysed data; C.I., K.E.D., J.A.V., V.A., K.A.B., Y.C., M.M.P., F.R., S.I.R., M.M., V.R.M., C.X.C., H.V. wrote the paper.

## Methods

### Culturing and nucleic acid extraction

*Ostreobium quekettii* (SAG culture collection strain 6.99, non-axenic) was cultured in F/2 media on a 14h/8h light/dark cycle at ~19°C. This strain was discovered growing on a culture of a small tropical marine red macroalga (*Acrothamnion preissii*) and isolated into culture. The strain is nested in a lineage of *Ostreobium* species found in scleractinian corals [61] and it readily colonises coral skeleton when it is provided as a substrate. This clearly shows that the strain is an appropriate representative for this limestone-burrowing coral-associated genus, despite it being initially isolated from a non-coral source. *Ostreobium* is known to produce flagellated spores and we think that a spore residing on the surface of the red alga is the most likely source of the strain. Total DNA was extracted using a modified cetyl trimethylammonium bromide (CTAB) method [62].

### Genome sequencing

We conducted three short-read sequencing runs using Illumina sequencing technology (Data S1) with 150 bp paired-end reads, for ~40 Gb data in total. For the first sequencing run, the total DNA was sheared to ~500 bp size fragments and processed using KAPA LTP Library Preparation Kit (Roche Sequencing Solutions, Pleasanton, California, USA) to prepare the DNA library, which was sequenced on the Illumina NextSeq 500 using PE 150 bp High Output Kit (Illumina, San Diego, California, USA), at Georgia Genomics and Bioinformatics Core (University of Georgia, USA). For the other two sequencing runs, the total DNA was sheared to ~350 bp fragments and the libraries were generated using TruSeq Nano DNA HT Sample preparation Kit (Illumina) following the manufacturer’s instructions. These DNA libraries were clustered on a cBot Cluster Generation System using HiSeq X HD PE Cluster Kit (Illumina) and sequenced on an Illumina HiSeq X Ten platform at Novogene, Beijing.

For long-reads Nanopore MinION sequencing (Oxford Nanopore Technologies), we used the Ligation Sequencing Kit 1D (SQK-LSK108) to prepare the DNA library, R9.5 chemistry, and Albacore v2.1.10 (https://github.com/Albacore/albacore) for basecalling.

### Assembly of genome and transcriptome data

Using the Illumina short-read data, an initial assessment of genome ploidy based on *k-* mer distribution was conducted using GenomeScope2 [63]. The result indicates that the data likely represent a diploid genome (*k* = 21; maximum 93.57% fit to the theoretical model). For genome assembly, we first combined all sequence data (Illumina short reads and Nanopore long reads) in a hybrid assembly using MaSuRCA v3.4.2 [64] at –ploidy 2 using cabog in the final step of assembling corrected mega-reads.

Transcriptome data of *Ostreobium*, assembled using Trinity v2.9.1 [65] in the *de novo* mode, were obtained from an earlier published work [66]. Using the genome assembly generated in this study, the transcriptome (RNA-Seq) reads (GenBank BioProject accession PRJEB35267) were assembled using Trinity in the “genome-guided” mode. Both assembled transcriptomes are used to guide identification of contaminant sequences and gene prediction (below).

### Identification and removal of contaminant sequences

We implemented a comprehensive strategy to systematically identify contaminants using GC content, read coverage, taxonomic annotations, and transcriptome data (Figure S6). Blobtools v1.1.1 [67] was first employed to generate a taxon-annotated GC-coverage plot. For each scaffold, the “bestsum” taxrule was applied based on results of BLASTn search (E ≤ 10-20) against genome sequences from all Bacteria, Archaea, viruses, Rhodophyta, Chlorophyta, Glaucophyta, and Chytridiomycota within the GenBank nucleotide (nt) database. The “bestsum” taxrule sums up all hits to a particular taxonomic group across the scaffold, ultimately designating the scaffold as originating from the taxon with the greatest overall alignment (bit) score. Coverage information for Blobtools was generated based on mapping of Illumina short reads and Nanopore long reads using Bowtie2 v2.3.5.1 [68] and Minimap2 v2.17 [69], respectively. Trinity transcripts (both *de novo* and genome-guided) were mapped against the genome assembly using Minimap2 (-ax splice -C5 --splice-flank=no --secondary=no) to identify the number of mapped transcripts and intron-containing transcripts for each genome scaffold.

We integrated this information in our strategy for identifying putative contaminant sequences using a decision tree (Figure S6). Briefly, for scaffolds that were not designated as chlorophyte sequences, we assessed the genome scaffold individually based on a combination of other criteria (i.e., taxonomic designation, outliers of GC-content and/or read coverage, and mapping of transcripts and intron-containing transcripts). Following Blobtools [67], for GC content and read coverage independently, we define an outlier as a value outside of the range of median ± interquartile range. For scaffolds that have no taxonomic designation, we do not exclude read-coverage outlier sequences if their GC is within the expected range, as these sequences likely represent repetitive regions of the genome. For scaffolds with a bacterial designation and mapped *Ostreobium* transcripts, we further require >10% of these transcripts to contain intron(s) to identify these scaffolds as non-contaminant. Full-length organellar genome sequences were also recovered using known *O. quekettii* plastid (KY509314.1) and mitochondrial (NC_045361.1) genomes as query in a BLASTn search and removed from the assembly. This process yielded the preliminary genome assembly (3134 scaffolds, N50 = 71.03Kb, total assembly size 151.9Mb).

### Genome size estimation

All the Illumina short reads from this study were mapped against the preliminary genome assembly with BWA v 0.7.17 [70] (mem mode; default parameters). Genome size was estimated based on *k*-mers of these reads that are free from putative contaminant sequences. The *k*-mers at *k* = 21 were enumerated using Jellyfish v2.3.0 [71], from which haploid genome size was estimated using Genomescope2.

### Generation of haploid genome assembly

Because the data were estimated to be diploid, using the preliminary, contaminant-free genome assembly, a haploid representation of the assembly was generated using purge_haplotigs [72] (-l 10 −m 55 −h 160 −j 101). This resulted in the final haploid genome assembly (2857 scaffolds, N50 = 73.43KB, total assembly size 146.263 MB), plus an additional 5.64Mbp of predicted heterozygous genomic regions.

### Ab initio prediction of protein-coding genes

We adapted the workflow from Chen et al. [73] for ab initio prediction of protein-coding genes in the haploid representation of *Ostreobium* genome assembly. This comprehensive workflow was initially designed for predicting genes from dinoflagellates, with modifications made to account for dinoflagellate-specific alternative splice sites; these modifications were ignored for predicting genes from *Ostreobium* because alternative intron splice sites are not expected in green algal genomes. Repetitive elements in the genome assembly were first predicted *de novo* using RepeatModeler v2.0.1 (http://www.repeatmasker.org/RepeatModeler/). These repeats were combined with known repeats in the RepeatMasker database (release 20181026) to generate a customised repeat library. All repetitive elements in the assembled genome scaffolds were then masked using RepeatMasker v4.1.0 (http://www.repeatmasker.org/) based on the customised repeat library, before they were subjected to prediction of genes.

The assembled transcripts were used as transcriptome evidence, and vector sequences were removed using SeqClean [74] based on UniVec (build 10) database. The PASA pipeline v2.4.1 [75] (--MAX_INTRON_LENGTH 70000 --ALIGNERS blat) and TransDecoder v5.5.0 [65] were first used to predict protein-coding genes (and the associated protein sequences) from the vector-trimmed transcriptome assemblies (hereinafter transcript-based genes). The predicted protein sequences from multi-exon transcript-based genes with complete 5′ and 3′-ends were searched (BLASTp, E ≤ 10^−20^) against RefSeq proteins (release 98). Genes with significant BLASTp hits (>80% query coverage) were retained. Transposable elements were identified using HHblits v3.1.0 [76] and Transposon-PSI (http://transposonpsi.sourceforge.net/), searching against the transposon subset of UniRef30 database (release 2020_03). Proteins putatively identified as transposable elements were removed. Those remaining were clustered using CD-HIT v4.8.1 (ID=75%) [77] to yield a non-redundant protein set, and the associated transcript-based genes were kept. These genes were further processed by the *Prepare_golden_genes_for_predictors.pl* script from the JAMg package (https://github.com/genomecuration/JAMg). This step yielded a set of high-quality “golden” genes, which were used as a training set for gene prediction using AUGUSTUS v3.3.3 [78] (allow_dss_consensus_gc=true and non_gt_dss_prob=1 for intron model; --softmasking=1 --gff3=on --UTR=on --exonnames=on) and SNAP [79] (at default setting). We also employed GeneMark-ES v4.48 [80] at default settings to generate predictions from the genome scaffolds, and MAKER protein2genome v2.31.10 [81] (at default setting) to make predictions based on homology to SwissProt proteins (downloaded 2 March 2020). We used unmasked repeat data for PASA, hard-masked repeats for GeneMark-ES and soft-marked repeat data for all other programs.

Subsequently, all genes predicted using GeneMark-ES, MAKER, PASA, SNAP and AUGUSTUS were integrated into a combined set using EvidenceModeler v1.1.1 [82], following a weighting scheme of GeneMark-ES 2, MAKER 8, PASA 10, SNAP 2, AUGUSTUS 6. The resulting EvidenceModeler predictions were retained if they were constructed using evidence from PASA, or using two or more other prediction methods.

Functional information of translated predicted genes was retrieved using search BLASTP against UniProt databases (Swiss-Prot and TrEMBL), KEGG’s annotation tool BlastKOALA [83]. Gene models of the genome dataset were annotated using InterProScan 5.39 [84] using InterPro (version 77.0 databases) of Pfam (32.0), SUPERFAMILY (1.75) and TIGRFAMs (15.0) [85]. The *Ostreobium* genomic data is available at https://doi.org/10.5281/zenodo.4012771.

### Orthogroups, phylogenetic analysis and BUSCO analysis

For comparative genomic analyses, we built a dataset containing genomes and gene annotations of 20 green algae and two land plants (Data S1). We used the OrthoFinder 2.3.7 [86] pipeline (default parameters) to cluster the potential orthologous protein families. Enrichment and depletion of domains in *Ostreobium* versus *Caulerpa lentillifera* were identified using Fisher’s exact tests with a false discovery rate correction (Benjamini-Hochberg FDR method) of 0.05. All statistical tests were carried out in R [87]. Subcellular localisation of sequences of interest was performed using PSORT [88], using the WoLF PSORT web server (https://wolfpsort.hgc.jp/) with the ‘plant’ option, and PredAlgo [89], using default settings.

We performed phylogenetic analyses for protein families relevant for photobiology and oxidative stress response, based on the orthogroups obtained from the OrthoFinder analysis described above. The protein sequences were aligned using PROMALS3D, with default parameters [90]. All phylogenetic trees were inferred using IQ-TREE 1.6.12 with the built-in model selection function, and branch support estimated using ultrafast bootstrap with 1,000 bootstrap replicates [91]. To identify the light-harvesting protein families associated with PSI (Lhca), we also included Lhca protein sequences from *Bryopsis corticulans* [24] and *Chlamydomonas reinhardtii* [23].

We ran BUSCO v4.1.4 [92] on the *Ostreobium* genomic data using the Eukaryota dataset (eukaryota_odb10.2020-09-10). This analysis identifies complete, duplicated, fragmented and missing genes that are expected to be present in a set of single-copy genes in the dataset. We ran identical BUSCO analyses on the *C. lentillifera, Ulva mutabilis* and *C. reinhardtii* genomes to allow direct comparison (Data S1).

### Metatranscriptome analysis of healthy and bleached corals

#### Experiment

*Orbicella faveolata* fragments (4 cm^2^) from three different colonies were collected in Petempiche Puerto Morelos, Quintana Roo, Mexico (N 20° 54′17.0″, W 86° 50′11.9″) at a depth between eight and nine meters in June 2013 (Permit registration MX-HR-010-MEX, Folio 036). The coral skeleton and live tissue were collected using a hammer and chisel and were transported in seawater to perform the experiment at the Instituto de Ciencias del Mar y Limnología, UNAM. Three fragments were placed in each control and “experimental” tank at ~28°C. Following 18 days of acclimation, heaters were turned on in the treatment tank to reach ~32°C. After five days of severe heat exposure, bleached corals were moved back into the control tank to recover at 28°C, where they remained for 38 days (post-bleaching period). Post-bleached and controlled coral fragments were flash-frozen and preserved in liquid nitrogen.

#### RNA library preparation and sequencing and data availability

Coral fragments were ground to a fine powder in liquid nitrogen. Total RNA was extracted using the mirVana miRNA Isolation Kit (Life Technologies) since this kit resulted in higher coral holobiont RNA yield and quality. RNA was purified and concentrated using the RNA Clean and Concentrator kit (Zymo Research, Irvine, USA). RNA quantification was assessed on a Nanodrop and Qubit using 2.0 RNA Broad Range Assay Kit (Invitrogen). Quality was verified using the Agilent Bioanalyzer 2100 (Agilent Technologies, Santa Clara, USA). Total RNA samples were sent for metatranscriptome sequencing to the US Department of Energy’s Joint Genome Institute (JGI), California. Samples were depleted in ribosomal RNA and enriched in mRNA from whole holobionts employing a cocktail of RiboZero kits (Epicenter), including the human/mouse/rat kit, plant kit and the mixed populations of gram-negative and gram-positive bacteria kit. Following the RiboZero protocol, mRNA was converted to cDNA and amplified. The libraries were sequenced on the Illumina HiSeq 2000 platform using 2 X 151 bp overlapping paired-end reads. Raw and filtered metatranscriptome sequence data, statistics and quality sequencing reports for the experiment are available at the US Department of Energy Joint Genome Institute (JGI)’s genome portal (https://genome.jgi.doe.gov/portal/) with accession codes 1086604, 1086606, 1086608, 1086610, 1086612, 1086614, Community Sequencing Project No. 1622).

#### Metatranscriptome analysis

The raw reads were quality-trimmed to Q10, adapter-trimmed and filtered for process artifacts using BBDuk [93]. Ribosomal RNA reads were removed by mapping against a trimmed version of the Silva database using BBmap (http://sourceforge.net/projects/bbmap). To generate a de novo reference metatranscriptome, cleaned reads from all samples per species (replicates from control and treatment fragments) were pooled and assembled using Trinity version 2.1.1 [65].

We performed a BLASTn search to identify and separate the different members of the holobiont using a local database. This database was built with the *Ostreobium* genome, the sea anemones *Nematostella vectensis* [94] and *Exaiptasia diaphana* (syn. *Aiptasia pallida*) [95], the corals *Acropora digitifera* [20] and *Orbicella faveolata* [96], and the Symbiodiniaceae genomes *Breviolum minutum* [97], *Fugacium kawagutii* [1] and *Symbiodinium microadriaticum* [98], as well as several *Breviolum* spp. transcriptomes [99]. The resulting *Ostreobium* transcriptomic portion (varied from 0.04% to 0.11% among the biological replicates) was used as a reference for gene quantification and subsequent differential gene expression analyses.

Kallisto [100] was used to perform a pseudoalignment and quantify transcript abundances, using the *Ostreobium* contigs derived from the metatranscriptome assembly as a reference. A comparison of counts per million, a correlation matrix and principal component analysis among samples was performed for a quality check of the replicates per species. We used three different biological replicates (i.e. coral colonies), which were split into three control samples and three post-bleaching treated samples for differential expression analysis performed using the DESeq2 software [101]. Differential Expressed Genes (DEGs) were defined by using a cutoff threshold of False Discovery Rate FDR <0.001 and log fold change of 2. Enzyme Commision numbers (EC) were retrieved from the Kyoto Encyclopedia of Genes and Genomes (KEGG) database using MEGAN5 [102] for genes from each holobiont member (i.e. host, Symbiodiniaceae and *Ostreobium*).

#### Taxonomic profile

To obtain the taxonomic profile of the metatranscriptome, quality control of the raw reads and assembled contigs was performed using Trim Galore v0.5 [103]. The KMA aligner v.1.3.2 [104] was used to map sequences against the NCBI nucleotide collection with the options’ 1t1 - mem_mode -and -apm p -ef’. CCMetagen v.1.2.5 [105] was used for an additional quality control and to obtain ranked taxonomic information. CCMetagen_merge was used with the options’-kr k −l Superkingdom-tlist Bacteria,Archaea’ to obtain the taxonomic information of the prokaryotic community.

## Supplemental Figure titles

**Figure S1. Maximum likelihood trees of light-harvesting complex proteins in green lineage. Related to Figure 2.**

(A) LHC associated with Photosystem I.

(B) LHC associated with Photosystem II.

Branch thickness shows the ultrafast bootstrap support results with 1000 replicates. *Ostreobium* proteins are in blue, *Caulerpa lentillifera* in teal, *Bryopsis corticulans* in purple, *Chlamydomonas reinhardtii* in orange and Streptophyta in green.

**Figure S2. Maximum likelihood tree of cryptochrome and photolyase photoreceptors. Related to Photobiology in a dark place subsection.**

Branch support results from an ultrafast bootstrap with 1000 replicates. *Ostreobium* proteins are in blue, *Caulerpa lentillifera* in teal, *Chlamydomonas reinhardtii* in orange and Streptophyta in green.

**Figure S3. Maximum likelihood tree of protein families related to oxidative stress response. Related to Figure 3.**

Branch thickness shows the support results from an ultrafast bootstrap with 1000 replicates. *Ostreobium* proteins are in blue, *Caulerpa lentillifera* in teal, *Chlamydomonas reinhardtii* in orange and *Ulva mutabilis* in purple. Superoxide dismutase, SOD; ascorbate peroxidase, APX; catalase, CAT; monodehydroascorbate reductase, MDHAR.

**Figure S4. The localisation of the reactions is based on data for C. reinhardtii []. Related to Life in an extreme environment section.**

Top and bottom organelles represent the mitochondrion and the chloroplast, respectively. The name of the genes encoding the relevant enzymes is shown in red and fermentation products are shown in purple boxes. Dashed lines represent transport across cellular compartments. The enzyme pyruvate decarboxylase (catalysing the production of acetaldehyde from pyruvate) was not found but the reaction is probably catalysed by another enzyme. We expect *Ostreobium* to use other electron acceptors to produce ATP and re-oxidize NAD(P)H and FADH2. *Ostreobium* possesses the enzymes required to produce succinate, lactate, formate, acetate, ethanol, alanine, and glycerol, but lacks H_2_ and acetate production from acetyl-CoA (PAT1/PAT2 and ACK1/ACK2). Several fermentation-related genes are present in multiple copies (Data S1), including two lactate dehydrogenases, four tandem copies of ALDH (aldehyde dehydrogenase), and six copies of malate dehydrogenase. The enzymes in the figure are: ADH1: alcohol dehydrogenase, ALAT: alanine aminotransferase, ALDH: aldehyde dehydrogenase, FDR: fumarate dehydrogenase, FUM: fumarase, GDP: glycerol-3-phosphate dehydrogenase, GPP: glycerol-3-phosphate phosphatase, LDH: lactate dehydrogenase, MDH: malate dehydrogenase, MME4: malic enzyme, PEPC: phosphoenolpyruvate carboxylase, PFL: pyruvate formate lyase, PPDK: pyruvate, phosphate dikinase.

**Figure S5. Comparative analysis of enriched and depleted InterPro domains in *Ostreobium* compared to the nonburrowing *Caulerpa lentillifera*.** Related to Life in an extreme environment section.

Significant differences relative to *C. lentillifera* (Fisher’s exact test, false discovery rate [FDR]-corrected p < 0.05). Z-scores represent the number of IPR hits normalized by the total number of hits per species. Grey numbers denote the total count of genes with the respective IPR domains in the genome selection.

**Figure S6. Systematical strategy to identify putative scaffolds from contaminants in the assembled genome. Related to STAR Methods (Identification and removal of contaminant sequences).**

## References

1. Lin, S., Cheng, S., Song, B., Zhong, X., Lin, X., Li, W., Li, L., Zhang, Y., Zhang, H., Ji, Z., et al. (2015). The Symbiodinium kawagutii genome illuminates dinoflagellate gene expression and coral symbiosis. Science 350, 691–694.

2. Robbins, S.J., Singleton, C.M., Chan, C.X., Messer, L.F., Geers, A.U., Ying, H., Baker, A., Bell, S.C., Morrow, K.M., Ragan, M.A., et al. (2019). A genomic view of the reef-building coral Porites lutea and its microbial symbionts. Nat Microbiol 4, 2090–2100.

3. Marcelino, V.R., Morrow, K.M., van Oppen, M.J.H., Bourne, D.G., and Verbruggen, H. (2017). Diversity and stability of coral endolithic microbial communities at a naturally high pCO2 reef. Mol Ecol 26, 5344–5357.

4. Ricci, F., Rossetto Marcelino, V., Blackall, L.L., Kuhl, M., Medina, M., and Verbruggen, H. (2019). Beneath the surface: community assembly and functions of the coral skeleton microbiome. Microbiome 7, 159.

5. Verbruggen, H., and Tribollet, A. (2011). Boring algae. Curr Biol 21, R876–877.

6. Tribollet, A. (2008). The boring microflora in modern coral reef ecosystems: a review of its roles. In Current Developments in Bioerosion. pp. 67–94.

7. Diaz-Pulido, G., and McCook, L.J. (2002). The fate of bleached corals: patterns and dynamics of algal recruitment. Marine Ecology Progress Series 232, 115–128.

8. Fine, M., Roff, G., Ainsworth, T.D., and Hoegh-Guldberg, O. (2006). Phototrophic microendoliths bloom during coral "white syndrome". Coral Reefs 25, 577–581.

9. Schlichter, D., Zscharnack, B., and Krisch, H. (1995). Transfer of photoassimilates from endolithic algae to coral tissue. Naturwissenschaften 82, 561–564.

10. Fine, M., and Loya, Y. (2002). Endolithic algae: an alternative source of photoassimilates during coral bleaching. Proc Biol Sci 269, 1205–1210.

11. Shashar, N., and Stambler, N. (1992). Endolithic algae within corals - life in an extreme environment. J. Exp. Mar. Biol. Ecol. 163, 277–286.

12. Magnusson, S.H., Fine, M., and Kuhl, M. (2007). Light microclimate of endolithic phototrophs in the scleractinian corals Montipora monasteriata and Porites cylindrica. Marine Ecology Progress Series 332, 119–128.

13. Koehne, B., Elli, G., Jennings, R.C., Wilhelm, C., and Trissl, H. (1999). Spectroscopic and molecular characterisation of a long wavelength absorbing antenna of Ostreobium sp. Biochim Biophys Acta 1412, 94–107.

14. Wilhelm, C., and Jakob, T. (2006). Uphill energy transfer from long-wavelength absorbing chlorophylls to PS II in Ostreobium sp. is functional in carbon assimilation. Photosynth Res 87, 323–329.

15. Kuhl, M., Holst, G., Larkum, A.W., and Ralph, P.J. (2008). Imaging of Oxygen Dynamics within the Endolithic Algal Community of the Massive Coral Porites Lobata(1). J Phycol 44, 541–550.

16. Leliaert, F., Smith, D.R., Moreau, H., Herron, M.D., Verbruggen, H., Delwiche, C.F., and De Clerck, O. (2012). Phylogeny and Molecular Evolution of the Green Algae. Critical Reviews in Plant Sciences 31, 1–46.

17. Verbruggen, H., Ashworth, M., LoDuca, S.T., Vlaeminck, C., Cocquyt, E., Sauvage, T., Zechman, F.W., Littler, D.S., Littler, M.M., Leliaert, F., et al. (2009). A multi-locus time-calibrated phylogeny of the siphonous green algae. Mol Phylogenet Evol 50, 642–653.

18. Tandon, K., Lu, C.Y., Chiang, P.W., Wada, N., Yang, S.H., Chan, Y.F., Chen, P.Y., Chang, H.Y., Chiou, Y.J., Chou, M.S., et al. (2020). Comparative genomics: Dominant coral-bacterium Endozoicomonas acroporae metabolises dimethylsulfoniopropionate (DMSP). ISME J 14, 1290–1303.

19. Kwong, W.K., Del Campo, J., Mathur, V., Vermeij, M.J.A., and Keeling, P.J. (2019). A widespread coral-infecting apicomplexan with chlorophyll biosynthesis genes. Nature 568, 103–107.

20. Shinzato, C., Shoguchi, E., Kawashima, T., Hamada, M., Hisata, K., Tanaka, M., Fujie, M., Fujiwara, M., Koyanagi, R., Ikuta, T., et al. (2011). Using the Acropora digitifera genome to understand coral responses to environmental change. Nature 476, 320–323.

21. Morosinotto, T., Breton, J., Bassi, R., and Croce, R. (2003). The nature of a chlorophyll ligand in Lhca proteins determines the far red fluorescence emission typical of photosystem I. J Biol Chem 278, 49223–49229.

22. Morosinotto, T., Castelletti, S., Breton, J., Bassi, R., and Croce, R. (2002). Mutation analysis of Lhca1 antenna complex. Low energy absorption forms originate from pigment-pigment interactions. J Biol Chem 277, 36253–36261.

23. Suga, M., Ozawa, S.I., Yoshida-Motomura, K., Akita, F., Miyazaki, N., and Takahashi, Y. (2019). Structure of the green algal photosystem I supercomplex with a decameric light-harvesting complex I. Nat Plants 5, 626–636.

24. Qin, X., Pi, X., Wang, W., Han, G., Zhu, L., Liu, M., Cheng, L., Shen, J.R., Kuang, T., and Sui, S.F. (2019). Structure of a green algal photosystem I in complex with a large number of light-harvesting complex I subunits. Nat Plants 5, 263–272.

25. Six, C., Worden, A.Z., Rodriguez, F., Moreau, H., and Partensky, F. (2005). New insights into the nature and phylogeny of prasinophyte antenna proteins: Ostreococcus tauri, a case study. Mol Biol Evol 22, 2217–2230.

26. Koziol, A.G., Borza, T., Ishida, K., Keeling, P., Lee, R.W., and Durnford, D.G. (2007). Tracing the evolution of the light-harvesting antennae in chlorophyll a/b-containing organisms. Plant Physiol 143, 1802–1816.

27. Neilson, J.A., and Durnford, D.G. (2010). Structural and functional diversification of the light-harvesting complexes in photosynthetic eukaryotes. Photosynth Res 106, 57–71.

28. Christa, G., Cruz, S., Jahns, P., de Vries, J., Cartaxana, P., Esteves, A.C., Serodio, J., and Gould, S.B. (2017). Photoprotection in a monophyletic branch of chlorophyte algae is independent of energy-dependent quenching (qE). New Phytol 214, 1132–1144.

29. Neilson, J.A., and Durnford, D.G. (2010). Evolutionary distribution of light-harvesting complex-like proteins in photosynthetic eukaryotes. Genome 53, 68–78.

30. Marcelino, V.R., Cremen, M.C., Jackson, C.J., Larkum, A.A., and Verbruggen, H. (2016). Evolutionary Dynamics of Chloroplast Genomes in Low Light: A Case Study of the Endolithic Green Alga Ostreobium quekettii. Genome Biol Evol 8, 2939–2951.

31. Rockwell, N.C., and Lagarias, J.C. (2020). Phytochrome evolution in 3D: deletion, duplication, and diversification. New Phytol 225, 2283–2300.

32. Duanmu, D., Bachy, C., Sudek, S., Wong, C.H., Jimenez, V., Rockwell, N.C., Martin, S.S., Ngan, C.Y., Reistetter, E.N., van Baren, M.J., et al. (2014). Marine algae and land plants share conserved phytochrome signaling systems. Proc Natl Acad Sci U S A 111, 15827–15832.

33. Kottke, T., Oldemeyer, S., Wenzel, S., Zou, Y., and Mittag, M. (2017). Cryptochrome photoreceptors in green algae: Unexpected versatility of mechanisms and functions. J Plant Physiol 217, 4–14.

34. Beel, B., Prager, K., Spexard, M., Sasso, S., Weiss, D., Muller, N., Heinnickel, M., Dewez, D., Ikoma, D., Grossman, A.R., et al. (2012). A flavin binding cryptochrome photoreceptor responds to both blue and red light in Chlamydomonas reinhardtii. Plant Cell 24, 2992–3008.

35. Duanmu, D., Rockwell, N.C., and Lagarias, J.C. (2017). Algal light sensing and photoacclimation in aquatic environments. Plant Cell Environ 40, 2558–2570.

36. Ogura, Y., Tokutomi, S., Wada, M., and Kiyosue, T. (2008). PAS/LOV proteins: A proposed new class of plant blue light receptor. Plant Signal Behav 3, 966–968.

37. Suzuki, J.Y., and Bauer, C.E. (1995). A prokaryotic origin for light-dependent chlorophyll biosynthesis of plants. Proc Natl Acad Sci U S A 92, 3749–3753.

38. Hunsperger, H.M., Randhawa, T., and Cattolico, R.A. (2015). Extensive horizontal gene transfer, duplication, and loss of chlorophyll synthesis genes in the algae. BMC Evol Biol 15, 16.

39. Cremen, M.C.M., Leliaert, F., Marcelino, V.R., and Verbruggen, H. (2018). Large Diversity of Nonstandard Genes and Dynamic Evolution of Chloroplast Genomes in Siphonous Green Algae (Bryopsidales, Chlorophyta). Genome Biol Evol 10, 1048–1061.

40. Yamazaki, S., Nomata, J., and Fujita, Y. (2006). Differential operation of dual protochlorophyllide reductases for chlorophyll biosynthesis in response to environmental oxygen levels in the cyanobacterium Leptolyngbya boryana. Plant Physiol 142, 911–922.

41. Gálová, E., Šalgovičová, I., Demko, V., Mikulová, K., Ševčovičová, A., Slováková, Ľ., Kyselá, V., and Hudák, J. (2008). A short overview of chlorophyll biosynthesis in algae. Biologia 63.

42. Verbruggen, H., Marcelino, V.R., Guiry, M.D., Cremen, M.C.M., and Jackson, C.J. (2017). Phylogenetic position of the coral symbiont Ostreobium (Ulvophyceae) inferred from chloroplast genome data. J Phycol 53, 790–803.

43. Marcelino, V.R., and Verbruggen, H. (2016). Multi-marker metabarcoding of coral skeletons reveals a rich microbiome and diverse evolutionary origins of endolithic algae. Sci Rep 6, 31508.

44. Del Cortona, A., Jackson, C.J., Bucchini, F., Van Bel, M., D’Hondt, S., Skaloud, P., Delwiche, C.F., Knoll, A.H., Raven, J.A., Verbruggen, H., et al. (2020). Neoproterozoic origin and multiple transitions to macroscopic growth in green seaweeds. Proc Natl Acad Sci U S A 117, 2551–2559.

45. Dullo, W.-C., Gektidis, M., Golubic, S., Heiss, G.A., Kampmann, H., Kiene, W., Kroll, D.K., Kuhrau, M.L., Radtke, G., Reijmer, J.G., et al. (1995). Factors controlling holocene reef growth: An interdisciplinary approach. Facies 32, 145–188.

46. Grossman, A.R., Bhaya, D., Apt, K.E., and Kehoe, D.M. (1995). Light-harvesting complexes in oxygenic photosynthesis: Diversity, Control, and Evolution. Annu Rev Genet 29, 231–288.

47. Foyer, C.H. (2018). Reactive oxygen species, oxidative signaling and the regulation of photosynthesis. Environ Exp Bot 154, 134–142.

48. Mallick, N., and Mohn, F.H. (2000). Reactive oxygen species: response of algal cells. Journal of Plant Physiology 157, 183–193.

49. Mao Che, L., Le Campion-Alsumard, T., Boury-Esnault, N., Payri, C., Golubic, S., and Bézac, C. (1996). Biodegradation of shells of the black pearl oyster, Pinctada margaritifera var. cumingii, by microborers and sponges of French Polynesia. Marine Biology 126, 509–519.

50. Bentis, C.J., Kaufman, L., and Golubic, S. (2000). Endolithic fungi in reef-building corals (Order : Scleractinia) are common, cosmopolitan, and potentially pathogenic. Biol Bull 198, 254–260.

51. Tribollet, A., Chauvin, A., and Cuet, P. (2019). Carbonate dissolution by reef microbial borers: a biogeological process producing alkalinity under different pCO2 conditions. Facies 65.

52. Garcia-Pichel, F., Ramirez-Reinat, E., and Gao, Q. (2010). Microbial excavation of solid carbonates powered by P-type ATPase-mediated transcellular Ca2+ transport. Proc Natl Acad Sci U S A 107, 21749–21754.

53. Machingura, M.C., Bajsa-Hirschel, J., Laborde, S.M., Schwartzenburg, J.B., Mukherjee, B., Mukherjee, A., Pollock, S.V., Forster, B., Price, G.D., and Moroney, J.V. (2017). Identification and characterisation of a solute carrier, CIA8, involved in inorganic carbon acclimation in Chlamydomonas reinhardtii. J Exp Bot 68, 3879–3890.

54. Matthews, J.L., Raina, J.B., Kahlke, T., Seymour, J.R., van Oppen, M.J.H., and Suggett, D.J. (2020). Symbiodiniaceae-bacteria interactions: rethinking metabolite exchange in reef-building corals as multi-partner metabolic networks. Environ Microbiol.

55. Croft, M.T., Lawrence, A.D., Raux-Deery, E., Warren, M.J., and Smith, A.G. (2005). Algae acquire vitamin B12 through a symbiotic relationship with bacteria. Nature 438, 90–93.

56. Helliwell, K.E., Wheeler, G.L., Leptos, K.C., Goldstein, R.E., and Smith, A.G. (2011). Insights into the evolution of vitamin B12 auxotrophy from sequenced algal genomes. Mol Biol Evol 28, 2921–2933.

57. Burriesci, M.S., Raab, T.K., and Pringle, J.R. (2012). Evidence that glucose is the major transferred metabolite in dinoflagellate-cnidarian symbiosis. J Exp Biol 215, 3467–3477.

58. Radecker, N., Pogoreutz, C., Voolstra, C.R., Wiedenmann, J., and Wild, C. (2015). Nitrogen cycling in corals: the key to understanding holobiont functioning? Trends Microbiol 23, 490–497.

59. Ferrer, L.M., and Szmant, A.M. (1988). Nutrient regeneration by the endolithic community in coral skeletons. Proc. 6th Coral Reef Symp. 3, 1–4.

60. Del Campo, J., Pombert, J.F., Slapeta, J., Larkum, A., and Keeling, P.J. (2017). The 'other' coral symbiont: Ostreobium diversity and distribution. ISME J 11, 296–299.

61. Gutner-Hoch, E., and Fine, M. (2011). Genotypic diversity and distribution of Ostreobium quekettii within scleractinian corals. Coral Reefs 30, 643–650.

62. Cremen, M.C.M., Huisman, J.M., Marcelino, V.R., and Verbruggen, H. (2016). Taxonomic revision of Halimeda (Bryopsidales, Chlorophyta) in south-western Australia. Australian Systematic Botany 29.

63. Ranallo-Benavidez, T.R., Jaron, K.S., and Schatz, M.C. (2020). GenomeScope 2.0 and Smudgeplot for reference-free profiling of polyploid genomes. Nat Commun 11, 1432.

64. Zimin, A.V., Puiu, D., Luo, M.C., Zhu, T., Koren, S., Marcais, G., Yorke, J.A., Dvorak, J., and Salzberg, S.L. (2017). Hybrid assembly of the large and highly repetitive genome of Aegilops tauschii, a progenitor of bread wheat, with the MaSuRCA mega-reads algorithm. Genome Res 27, 787–792.

65. Haas, B.J., Papanicolaou, A., Yassour, M., Grabherr, M., Blood, P.D., Bowden, J., Couger, M.B., Eccles, D., Li, B., Lieber, M., et al. (2013). De novo transcript sequence reconstruction from RNA-seq using the Trinity platform for reference generation and analysis. Nat Protoc 8, 1494–1512.

66. Repetti, S.I., Jackson, C.J., Judd, L.M., Wick, R.R., Holt, K.E., and Verbruggen, H. (2020). The inflated mitochondrial genomes of siphonous green algae reflect processes driving expansion of noncoding DNA and proliferation of introns. PeerJ 8, e8273.

67. Laetsch, D.R., and Blaxter, M.L. (2017). BlobTools: Interrogation of genome assemblies. F1000Research 6.

68. Langmead, B., and Salzberg, S.L. (2012). Fast gapped-read alignment with Bowtie 2. Nat Methods 9, 357–359.

69. Li, H. (2018). Minimap2: pairwise alignment for nucleotide sequences. Bioinformatics 34, 3094–3100.

70. Li, H., and Durbin, R. (2009). Fast and accurate short read alignment with Burrows-Wheeler transform. Bioinformatics 25, 1754–1760.

71. Marcais, G., and Kingsford, C. (2011). A fast, lock-free approach for efficient parallel counting of occurrences of k-mers. Bioinformatics 27, 764–770.

72. Roach, M.J., Schmidt, S.A., and Borneman, A.R. (2018). Purge Haplotigs: allelic contig reassignment for third-gen diploid genome assemblies. BMC Bioinformatics 19, 460.

73. Chen, Y., Gonzalez-Pech, R.A., Stephens, T.G., Bhattacharya, D., and Chan, C.X. (2020). Evidence That Inconsistent Gene Prediction Can Mislead Analysis of Dinoflagellate Genomes. J Phycol 56, 6–10.

74. Chen, Y.A., Lin, C.C., Wang, C.D., Wu, H.B., and Hwang, P.I. (2007). An optimised procedure greatly improves EST vector contamination removal. BMC Genomics 8, 416.

75. Haas, B.J., Delcher, A.L., Mount, S.M., Wortman, J.R., Smith, R.K., Jr., Hannick, L.I., Maiti, R., Ronning, C.M., Rusch, D.B., Town, C.D., et al. (2003). Improving the Arabidopsis genome annotation using maximal transcript alignment assemblies. Nucleic Acids Res 31, 5654–5666.

76. Remmert, M., Biegert, A., Hauser, A., and Soding, J. (2011). HHblits: lightning-fast iterative protein sequence searching by HMM-HMM alignment. Nat Methods 9, 173–175.

77. Li, W., and Godzik, A. (2006). Cd-hit: a fast program for clustering and comparing large sets of protein or nucleotide sequences. Bioinformatics 22, 1658–1659.

78. Stanke, M., Keller, O., Gunduz, I., Hayes, A., Waack, S., and Morgenstern, B. (2006). AUGUSTUS: ab initio prediction of alternative transcripts. Nucleic Acids Res 34, W435–439.

79. Korf, I. (2004). Gene finding in novel genomes. BMC Bioinformatics 5, 59.

80. Lomsadze, A., Gemayel, K., Tang, S., and Borodovsky, M. (2018). Modeling leaderless transcription and atypical genes results in more accurate gene prediction in prokaryotes. Genome Res 28, 1079–1089.

81. Holt, C., and Yandell, M. (2011). MAKER2: an annotation pipeline and genome-database management tool for second-generation genome projects. BMC Bioinformatics 12, 491.

82. Haas, B.J., Salzberg, S.L., Zhu, W., Pertea, M., Allen, J.E., Orvis, J., White, O., Buell, C.R., and Wortman, J.R. (2008). Automated eukaryotic gene structure annotation using EVidenceModeler and the Program to Assemble Spliced Alignments. Genome Biol 9, R7.

83. Kanehisa, M., Sato, Y., and Morishima, K. (2016). BlastKOALA and GhostKOALA: KEGG Tools for Functional Characterisation of Genome and Metagenome Sequences. J Mol Biol 428, 726–731.

84. Jones, P., Binns, D., Chang, H.Y., Fraser, M., Li, W., McAnulla, C., McWilliam, H., Maslen, J., Mitchell, A., Nuka, G., et al. (2014). InterProScan 5: genome-scale protein function classification. Bioinformatics 30, 1236–1240.

85. Mitchell, A.L., Attwood, T.K., Babbitt, P.C., Blum, M., Bork, P., Bridge, A., Brown, S.D., Chang, H.Y., El-Gebali, S., Fraser, M.I., et al. (2019). InterPro in 2019: improving coverage, classification and access to protein sequence annotations. Nucleic Acids Res 47, D351–D360.

86. Emms, D.M., and Kelly, S. (2019). OrthoFinder: phylogenetic orthology inference for comparative genomics. Genome Biol 20, 238.

87. Team, R.D.C. (2018). R: A Language and Environment for Statistical Computing. (Vienna, Austria: R Foundation for Statistical Computing).

88. Horton, P., Park, K.J., Obayashi, T., Fujita, N., Harada, H., Adams-Collier, C.J., and Nakai, K. (2007). WoLF PSORT: protein localisation predictor. Nucleic Acids Res 35, W585–587.

89. Tardif, M., Atteia, A., Specht, M., Cogne, G., Rolland, N., Brugiere, S., Hippler, M., Ferro, M., Bruley, C., Peltier, G., et al. (2012). PredAlgo: a new subcellular localisation prediction tool dedicated to green algae. Mol Biol Evol 29, 3625–3639.

90. Pei, J., Kim, B.H., and Grishin, N.V. (2008). PROMALS3D: a tool for multiple protein sequence and structure alignments. Nucleic Acids Res 36, 2295–2300.

91. Nguyen, L.T., Schmidt, H.A., von Haeseler, A., and Minh, B.Q. (2015). IQ-TREE: a fast and effective stochastic algorithm for estimating maximum-likelihood phylogenies. Mol Biol Evol 32, 268–274.

92. Waterhouse, R.M., Seppey, M., Simao, F.A., Manni, M., Ioannidis, P., Klioutchnikov, G., Kriventseva, E.V., and Zdobnov, E.M. (2018). BUSCO Applications from Quality Assessments to Gene Prediction and Phylogenomics. Mol Biol Evol 35, 543–548.

93. Bushnell, B. (2019). Bbtools Software Package. Volume 2019.

94. Putnam, N.H., Srivastava, M., Hellsten, U., Dirks, B., Chapman, J., Salamov, A., Terry, A., Shapiro, H., Lindquist, E., Kapitonov, V.V., et al. (2007). Sea anemone genome reveals ancestral eumetazoan gene repertoire and genomic organisation. Science 317, 86–94.

95. Baumgarten, S., Simakov, O., Esherick, L.Y., Liew, Y.J., Lehnert, E.M., Michell, C.T., Li, Y., Hambleton, E.A., Guse, A., Oates, M.E., et al. (2015). The genome of Aiptasia, a sea anemone model for coral symbiosis. Proc Natl Acad Sci U S A 112, 11893–11898.

96. Prada, C., Hanna, B., Budd, A.F., Woodley, C.M., Schmutz, J., Grimwood, J., Iglesias-Prieto, R., Pandolfi, J.M., Levitan, D., Johnson, K.G., et al. (2016). Empty Niches after Extinctions Increase Population Sizes of Modern Corals. Curr Biol 26, 3190–3194.

97. Shoguchi, E., Shinzato, C., Kawashima, T., Gyoja, F., Mungpakdee, S., Koyanagi, R., Takeuchi, T., Hisata, K., Tanaka, M., Fujiwara, M., et al. (2013). Draft assembly of the Symbiodinium minutum nuclear genome reveals dinoflagellate gene structure. Curr Biol 23, 1399–1408.

98. Aranda, M., Li, Y., Liew, Y.J., Baumgarten, S., Simakov, O., Wilson, M.C., Piel, J., Ashoor, H., Bougouffa, S., Bajic, V.B., et al. (2016). Genomes of coral dinoflagellate symbionts highlight evolutionary adaptations conducive to a symbiotic lifestyle. Sci Rep 6, 39734.

99. Parkinson, J.E., Baumgarten, S., Michell, C.T., Baums, I.B., LaJeunesse, T.C., and Voolstra, C.R. (2016). Gene Expression Variation Resolves Species and Individual Strains among Coral-Associated Dinoflagellates within the Genus Symbiodinium. Genome Biol Evol 8, 665–680.

100. Bray, N.L., Pimentel, H., Melsted, P., and Pachter, L. (2016). Near-optimal probabilistic RNA-seq quantification. Nature Biotechnology 34, 525–527.

101. Love, M.I., Huber, W., and Anders, S. (2014). Moderated estimation of fold change and dispersion for RNA-seq data with DESeq2. Genome Biol 15, 550.

102. Huson, D.H., Beier, S., Flade, I., Gorska, A., El-Hadidi, M., Mitra, S., Ruscheweyh, H.J., and Tappu, R. (2016). MEGAN Community Edition - Interactive Exploration and Analysis of Large-Scale Microbiome Sequencing Data. PLoS Comput Biol 12, e1004957.

103. Krueger, F. (2015). Trim Galore: A wrapper tool around Cutadapt and FastQC to consistently apply quality and adapter trimming to FastQ files, with some extra functionality for MspI-digested RRBS-type (Reduced Representation Bisufite-Seq) libraries. (http://www.bioinformatics.babraham.ac.uk/projects/trim_galore/).

104. Clausen, P., Aarestrup, F.M., and Lund, O. (2018). Rapid and precise alignment of raw reads against redundant databases with KMA. BMC Bioinformatics 19, 307.

105. Marcelino, V.R., Clausen, P., Buchmann, J.P., Wille, M., Iredell, J.R., Meyer, W., Lund, O., Sorrell, T.C., and Holmes, E.C. (2020). CCMetagen: comprehensive and accurate identification of eukaryotes and prokaryotes in metagenomic data. Genome Biol 21, 103.

106. Catalanotti, C., Yang, W., Posewitz, M.C., and Grossman, A.R. (2013). Fermentation metabolism and its evolution in algae. Front Plant Sci 4, 150.

