## Supplemental figures for "Genomic adaptations to an endolithic lifestyle in the coral-associated alga *Ostreobium*"

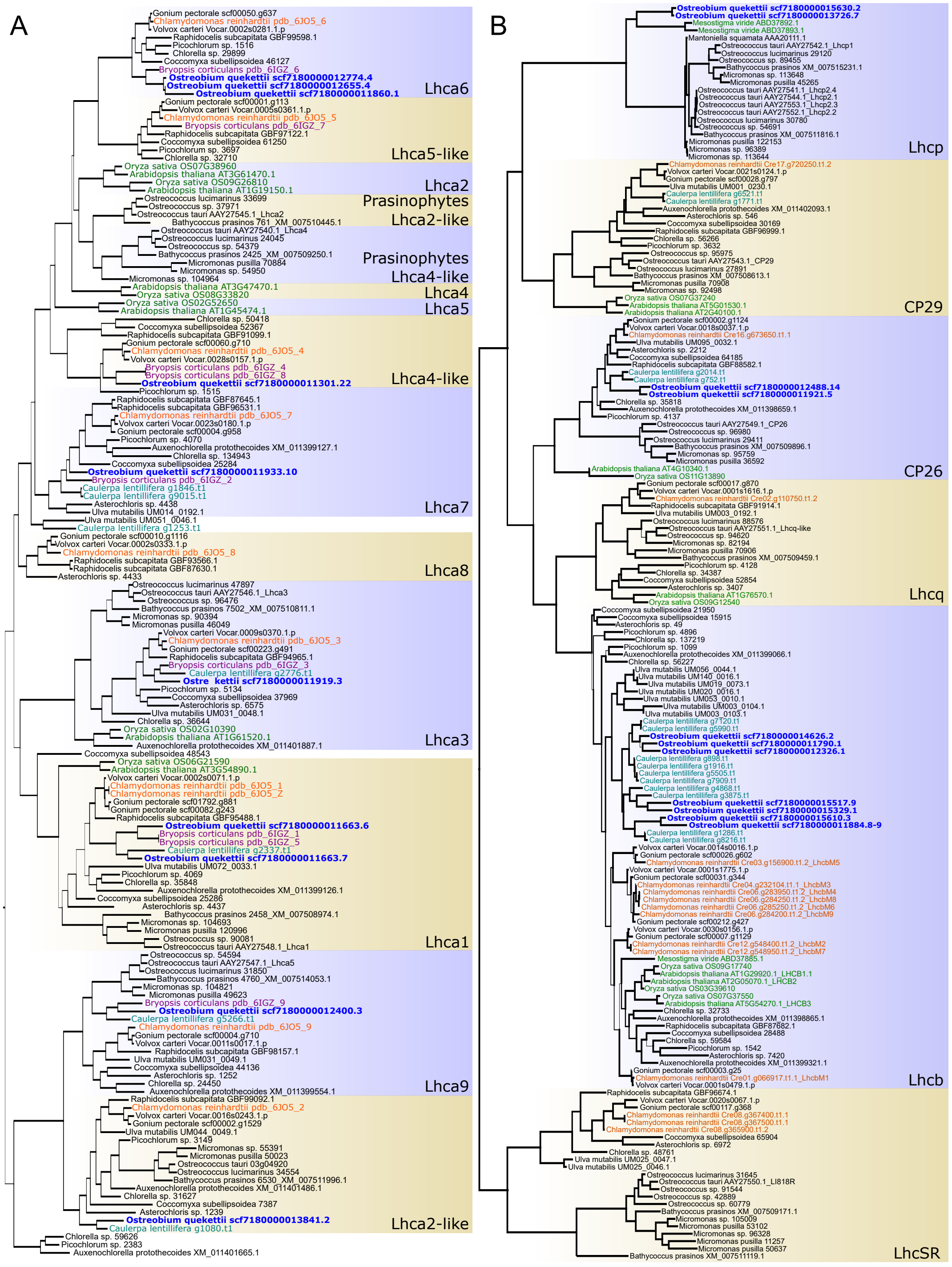


**Figure S1. Maximum likelihood trees of light-harvesting complex proteins in green lineage. Related to Figure 2**

(A) LHC associated with Photosystem I.

(B) LHC associated with Photosystem II.

Branch thickness shows the ultrafast bootstrap support results with 1000 replicates. *Ostreobium* proteins are in blue, *Caulerpa lentillifera* in teal, *Bryopsis corticulans* in purple, *Chlamydomonas reinhardtii* in orange and Streptophyta in green.

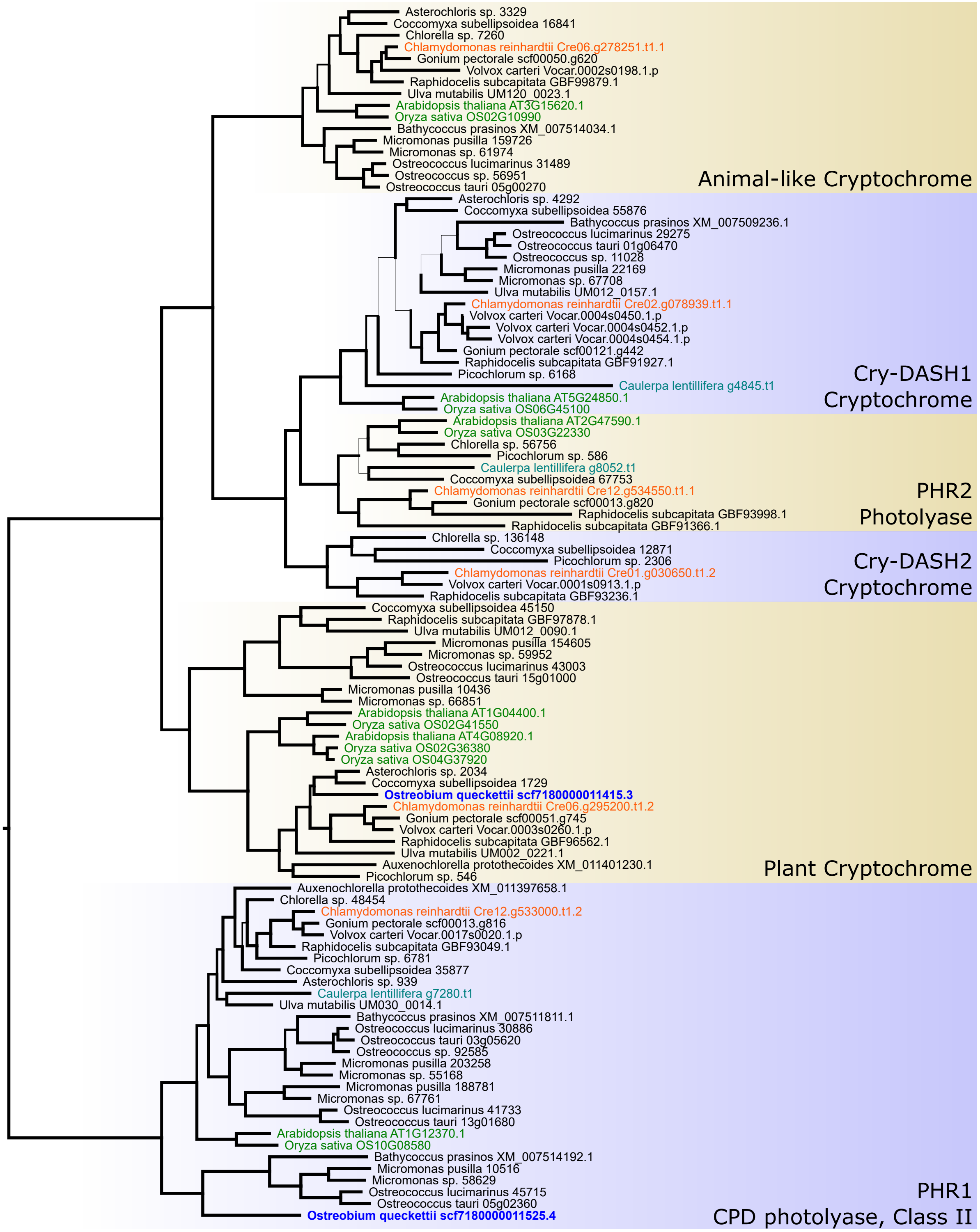


**Figure S2. Maximum likelihood tree of cryptochrome and photolyase photoreceptors. Related to *Photobiology in a dark place* subsection.**
Branch support results from an ultrafast bootstrap with 1000 replicates. *Ostreobium* proteins are in blue, *Caulerpa lentillifera* in teal, *Chlamydomonas reinhardtii* in orange and Streptophyta in green.

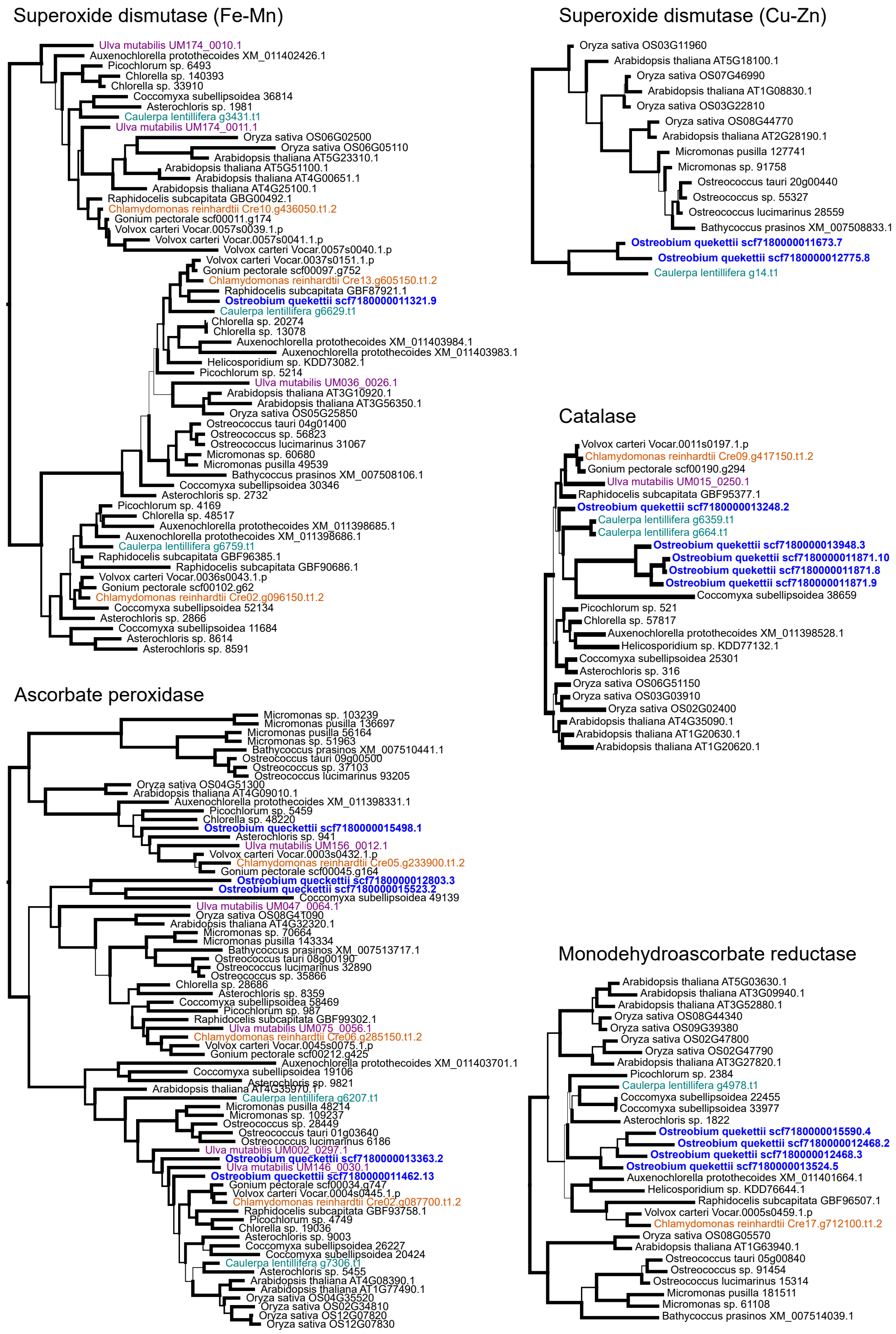


**Figure S3. Maximum likelihood tree of protein families related to oxidative stress response. Related to Figure 3.** Branch thickness shows the support results from an ultrafast bootstrap with 1000 replicates. *Ostreobium* proteins are in blue, *Caulerpa lentillifera* in teal, *Chlamydomonas reinhardtii* in orange and *Ulva mutabilis* in purple. Superoxide dismutase, SOD; ascorbate peroxidase, APX; catalase, CAT; monodehydroascorbate reductase, MDHAR.

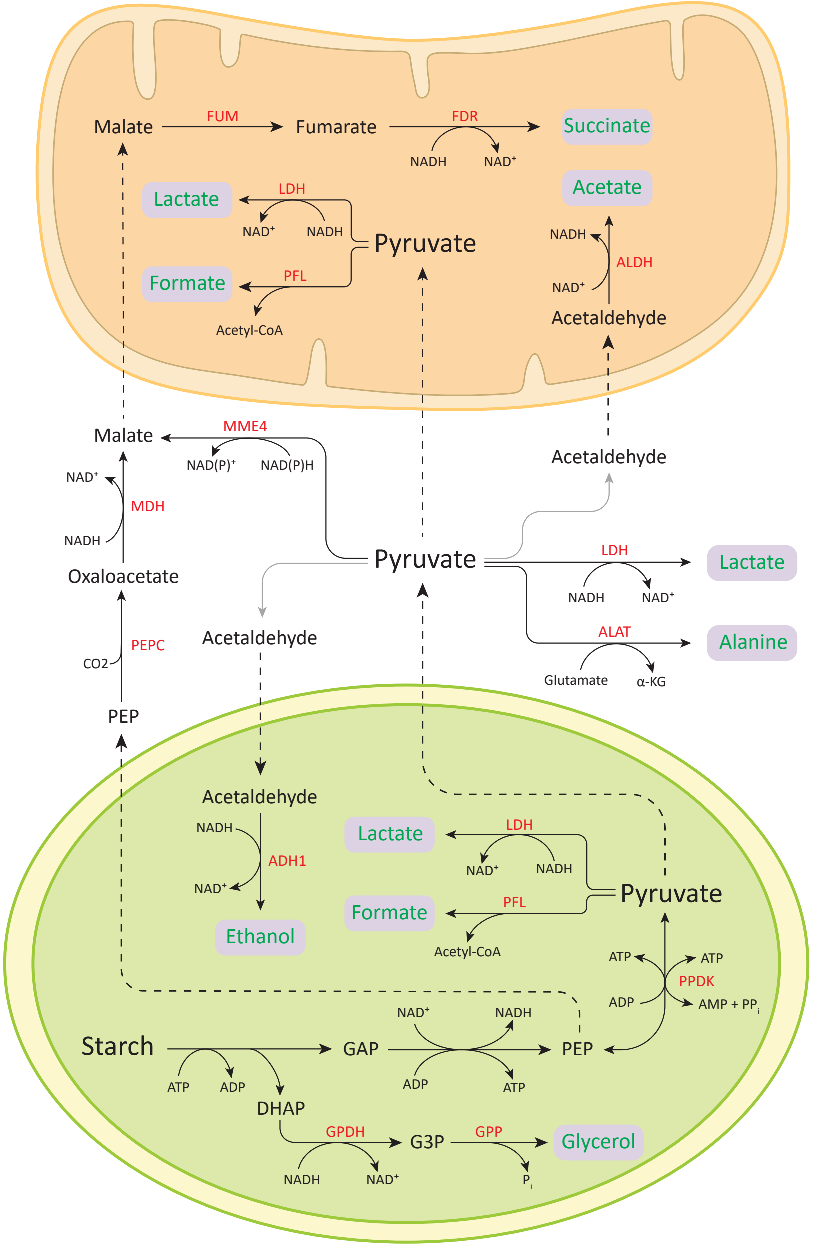


**Figure S4. The localisation of the reactions is based on data for *C. reinhardtii* (1). Related to *Life in an extreme environment* section.**
Top and bottom organelles represent the mitochondrion and the chloroplast, respectively. The name of the genes encoding the relevant enzymes is shown in red and fermentation products are shown in purple boxes. Dashed lines represent transport across cellular compartments. The enzyme pyruvate decarboxylase (catalysing the production of acetaldehyde from pyruvate) was not found but the reaction is probably catalysed by another enzyme. We expect *Ostreobium* to use other electron acceptors to produce ATP and re-oxidize NAD(P)H and FADH2. *Ostreobium* possesses the enzymes required to produce succinate, lactate, formate, acetate, ethanol, alanine, and glycerol, but lacks $H^2$ and acetate production from acetyl-CoA (PAT1/PAT2 and ACK1/ACK2). Several fermentation-related genes are present in multiple copies (Data S1), including two lactate dehydrogenases, four tandem copies of ALDH (aldehyde dehydrogenase), and six copies of malate dehydrogenase. The enzymes in the figure are: ADH1: alcohol dehydrogenase, ALAT: alanine aminotransferase, ALDH: aldehyde dehydrogenase, FDR: fumarate dehydrogenase, FUM: fumarase, GDP: glycerol-3-phosphate dehydrogenase, GPP: glycerol-3-phosphate phosphatase, LDH: lactate dehydrogenase, MDH: malate dehydrogenase, MME4: malic enzyme, PEPC: phosphoenolpyruvate carboxylase, PFL: pyruvate formate lyase, PPDK: pyruvate, phosphate dikinase.

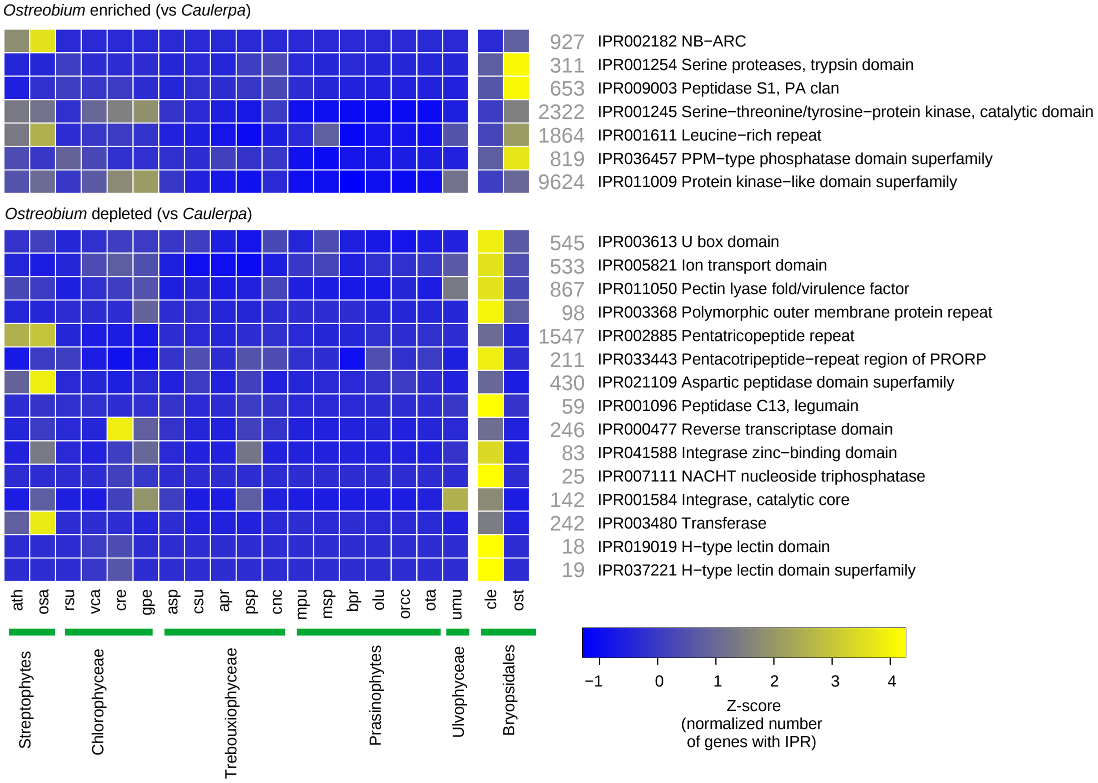


**Figure S5. Comparative analysis of enriched and depleted InterPro domains in *Ostreobium* compared to the non-burrowing *Caulerpa lentillifera*. Related to *Life in an extreme environment* section.**

Significant differences relative to *C. lentillifera* (Fisher's exact test, false discovery rate [FDR]-corrected $p < 0.05$). Z-scores represent the number of IPR hits normalized by the total number of hits per species. Grey numbers denote the total count of genes with the respective IPR domains in the genome selection.

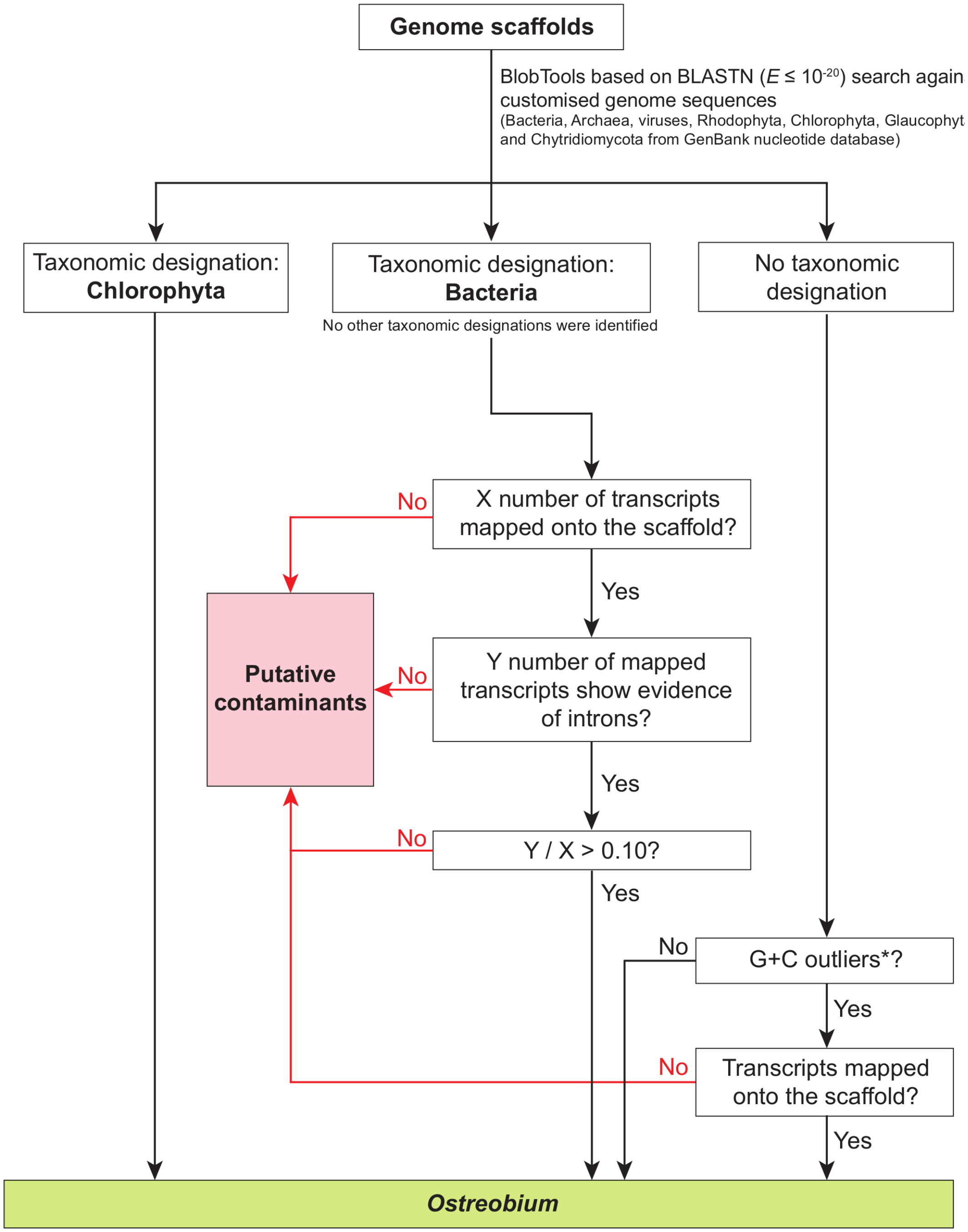


*: outliers are determined using BLOBtools. Scaffolds for which G+C content is external to the range of median ± 1.5 × interquartile range (IQR) are considered as outliers.

**Figure S6. Systematical strategy to identify putative scaffolds from contaminants in the assembled genome. Related to STAR Methods**

### Supplemental Reference

1. Catalanotti, C., Yang, W., Posewitz, M.C., and Grossman, A.R. (2013). Fermentation metabolism and its evolution in algae. Front Plant Sci *4*, 150.
